# Heparan sulfate selectively inhibits the collagenase activity of matrix metalloproteinase 13

**DOI:** 10.64898/2026.08.21.746339

**Authors:** Huanmeng Hao, Guowei Su, Jian Liu, Ding Xu

## Abstract

Matrix metalloproteinase 13 (MMP13) is a zinc-dependent protease that plays key roles in extracellular matrix remodeling. Like several other MMPs, MMP13 has been shown to interact with heparan sulfate (HS), a highly sulfated glycosaminoglycan found at the cell surface and in the extracellular matrix, but the significance of the interaction remains unknown. Here we report that while zymogen and mature forms of MMP13 both bind HS with high affinity, their interactions with HS display markedly different characteristics in terms of preferred HS structure and binding kinetics. By structure-guided mutagenesis, we identified a large HS-binding site of MMP13 consists of 10 residues in the hemopexin domain, 3 residues in the catalytic domain, and 2 residues in the linker region. While these basic residues participate in binding to both zymogen and mature forms of MMP13, the relative contribution of many residues differs substantially between the two forms, which likely contributes to their distinct HS-binding characteristics. Binding of HS to mature MMP13 resulted in selective inhibition of the collagenase activity of MMP13 in a length- and sulfation-dependent manner, but the binding had no effect on degradation of non-collagen substrates. Mechanistically, the inhibitory effect of HS likely results from reduced interdomain flexibility after binding of HS, and/or HS-induced dimerization of MMP13. In sum, our study establishes HS as a multifaceted regulator of MMP13 activity, and discovers that the HS-binding site of MMP13 is a novel exosite that can be targeted to inhibits its collagenase activity.

## INTRODUCTION

Matrix metalloproteinases (MMPs) are zinc-dependent endopeptidases that primarily function in degrading numerous extracellular matrix (ECM) substrates, including collagens, proteoglycans, and glycoproteins, thereby contributing to ECM remodeling^1–6^. MMPs also regulate diverse signaling pathways by directly cleaving signaling molecules or generating bioactive fragments^2,3^. Therefore, the MMPs are essential players in many physiological processes, such as embryonic development, tissue homeostasis, wound repair, and inflammation^7–9^. Accordingly, their biological functions are tightly regulated at multiple levels: gene expression, zymogen activation, localization and inactivation by tissue inhibitors of metalloproteinases (TIMPs)^10–12^. Dysregulated MMP activity thereby disrupts the balance in matrix synthesis and degradation, resulting in pathological matrix breakdown in pathological conditions such as osteoarthritis (OA), cancer, fibrosis, and inflammatory diseases^9,13,14^.

MMP1, MMP8, and MMP13 are three classical collagenases in MMP superfamily, capable of cleaving intact fibrillar collagens^15^. Among them, MMP13, also known as collagenase 3, is particularly robust in degrading type I and type II collagens, which are the major components in bone and articular cartilage, respectively^14,16,17^. The critical role of MMP13 in skeletal homeostasis is demonstrated by human genetic disorders such as spondyloepimetaphyseal dysplasia and metaphyseal anadysplasia^18^. These disorders are characterized by growth plate abnormity and shortening of limbs, which are caused by missense mutations of *MMP13*^19^. Key skeletal phenotypes of these human disorders are nicely recapitulated by *Mmp13*-null mice^20–22^. MMP13 is mainly expressed by chondrocytes in cartilage and osteoblasts in bone^23^. In a healthy joint, chondrocytes constitutively express and secret MMP13, but its extracellular concentration is limited by rapid LRP1-mediated endocytosis and lysosomal clearance^24^. Inflammatory cytokines, such as IL-1β and TNFα, induce expression of MMP13 robustly and exacerbate the degradation of cartilage, which is a hallmark in OA pathology^16,25–29^.

Heparan sulfate (HS) is a highly sulfated glycosaminoglycan (GAG) that is ubiquitously present in the ECM^30,31^. Previous studies have identified alterations in HS abundance and structure in OA cartilage and shown that manipulating HS structure can affect OA progression in animal models^32–35^. There are many indications that HS interacts with MMPs to regulate their compartmentalization, activation, activity, and internalization^24,36–46^. For instance, cell surface HS serves as extracellular docking sites for multiple MMPs, such as MMP7 and MMP9, and may concentrate MMPs near their substrates^36,43^. Highly sulfated GAGs, including heparin, were also found to accelerate pro-MMP7 activation and increase its activity toward selected substrates^40,44,45^. Importantly, MMP13 is known to bind heparin, and heparin binding was reported to inhibit LRP1-mediated internalization of MMP13^24^. Thus, an interaction between HS and MMP13 might be physiologically relevant and add an additional layer of modulation to MMP13 biology. However, the physiological significance of HS-MMP13 interactions, and the molecular basis and functional consequences of this interaction remains unknown.

In this study, we investigated the structural details of HS-MMP13 interaction and determined the functional impact of HS on MMP13 activity. We discovered that proMMP13 and mature MMP13 showed distinct HS binding kinetics and structural preferences. HS oligosaccharide also induces dimerization of mature MMP13 but not proMMP13. We further mapped the HS-binding residues and found overlapping but distinct binding surfaces in proMMP13 and mature MMP13. Importantly, we found that HS could selectively inhibit the collagenase activity of MMP13 in a sulfation- and length-dependent manner. Our findings strongly suggest HS can function as a selective regulator of MMP13 collagenase activity. This insight identifies the HS-binding surface of MMP13 as a novel structural element of MMP13, which might be targeted to develop novel selective MMP13 inhibitors.

## RESULTS

### proMMP13 and mature MMP13 exhibit distinct binding characteristics to HS

To examine the HS-binding properties of MMP13, we expressed human pro-MMP13 in Expi293 cells. As reported previously, we found secreted proMMP13 binds to Heparin-Sepharose column, and elution of proMMP13 from heparin column requires a salt concentration of 690mM (Fig. 1A). Processing of proMMP13 by trypsin generates mature MMP13 after removal of the propeptide (Fig. 1B). We found mature MMP13 binds to the heparin-affinity column even stronger, and elution from the column requires 770mM of NaCl (Fig. 1A).

**Figure 1.**
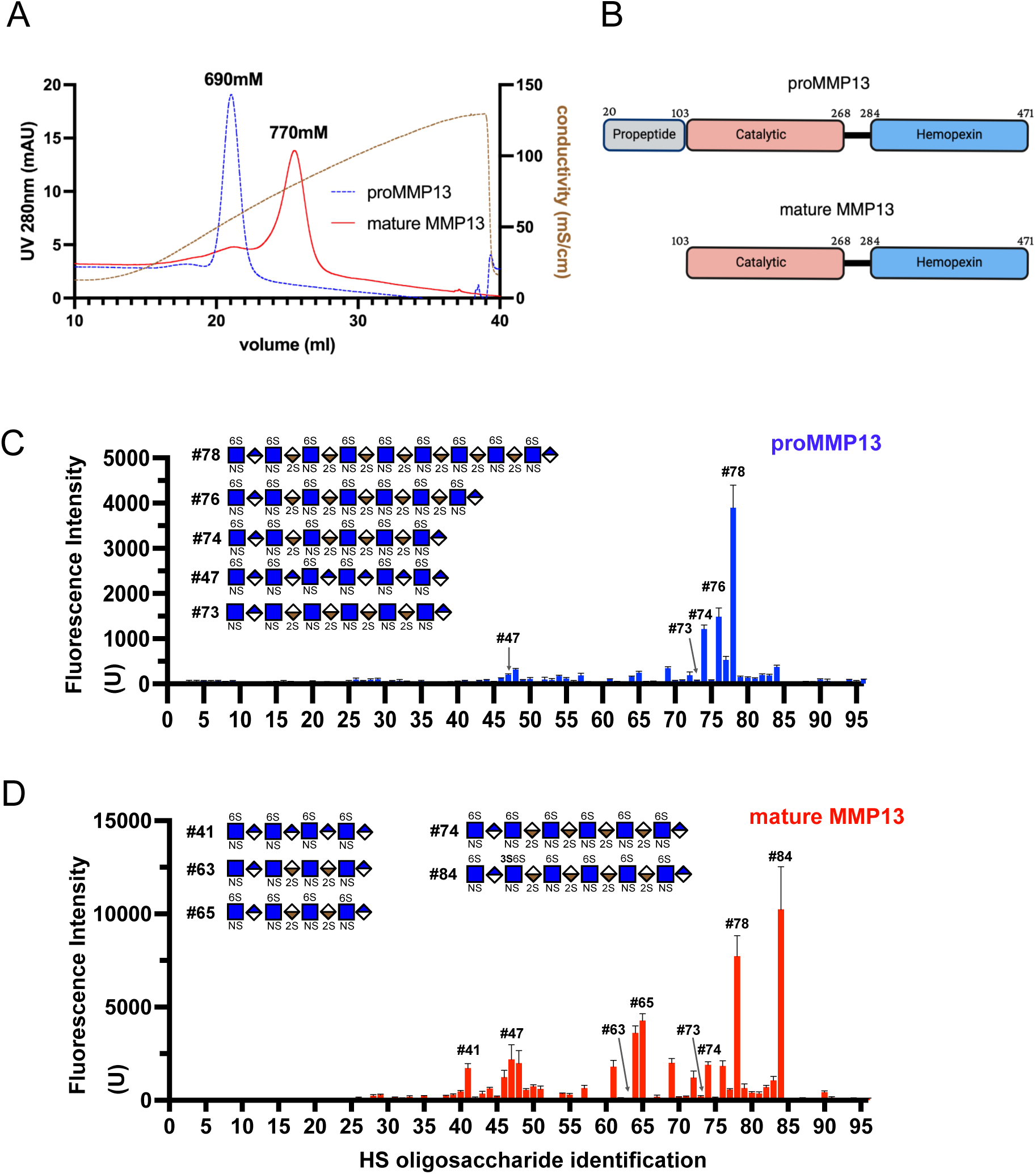
Pro-MMP13 and mature MMP13 exhibit distinct binding preferences to HS. (A) Heparin-Sepharose column elution profiles of recombinant human proMMP13 and mature MMP13. The dashed brown line indicates the linear NaCl gradient from 150 mM to 2 M, measured by conductivity, mS/cm. (B) Schematic view of the domain structure of MMP13. The linker region is shown in thick black line. Domain junctions are labeled by the residue number. (C, D) HS oligosaccharide microarray analysis of proMMP13 (C) and mature MMP13 binding (D). Structures of representative oligosaccharides are shown. Data are presented as mean SD of 36 printed spots for each oligosaccharide. This list of all oligosaccharides included in the microarray is shown in Supplemental Figure. 1.

Next we determined the structural requirements of HS oligosaccharides for MMP13 binding using a glycan microarray consisting of 96 structurally defined HS oligosaccharides with various lengths and sulfation patterns. proMMP13 only binds to a very limited set of HS oligosaccharides (compound 74, 76 and 78) with the minimum length required for pro-MMP13 binding is 12mer (compound 74) (Fig. 1C). In addition, fully sulfated 12mer appears to be required for binding as under-sulfated 12mer (compound 47 and 73) showed no binding. In contrast, mature MMP13 shows a broader binding spectrum, and HS 8mer is the minimum length required for binding (compound 41 and 65) (Fig. 1D). Interestingly, we found under-sulfated HS oligosaccharides with only NS and 6S groups (compound 41 and 47) can still bind well to mature MMP13, while HS oligosaccharides with only NS and 2S groups (compound 63 and 73) display no binding at all. This result suggests that 6-O-sulfation is required, while 2-O-sulfation is dispensable for binding to mature MMP13. We also found that an additional 3-O-sulfation strikingly increases the mature MMP13 binding signal (compound 84 compared to 74) (Fig. 1D), suggesting that 3-O-sulfation greatly promotes mature MMP13-HS interaction.

Using biolayer interferometry analysis, we determined the kinetic binding affinity of proMMP13 and mature MMP13 to immobilized HS 12mer (compound 74). Despite similar kinetic binding affinity of mature MMP13 and proMMP13 (42 and 57 nM, respectively), BLI assay revealed that the binding characteristics of proMMP13 and mature MMP13 are completely different (Fig. 2). proMMP13 displays a 5-fold faster association rate (6.6 x 10^4^ 1/Ms) compared to mature MMP13 (1.2 x 10^4^ 1/Ms). But the dissociation rate of proMMP13 is also 8-fold faster (3.8 x 10^-3^ 1/s) than the mature MMP13 (5 x 10^-4^ 1/s).

**Figure 2.**
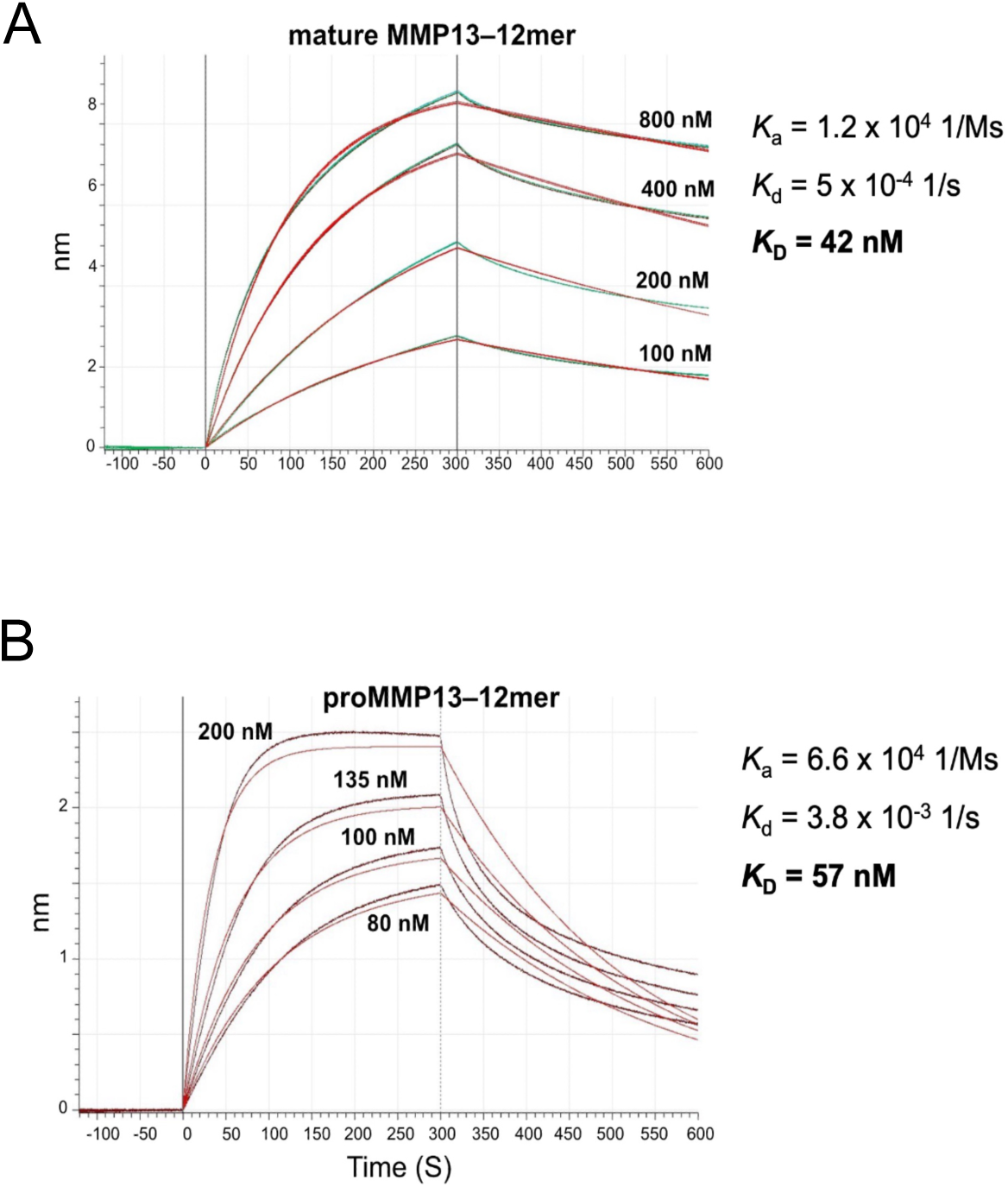
Biolayer interferometry kinetic analysis of binding of proMMP13 and mature MMP13 to 12mer. Biotinylated HS 12mer (#74 in Fig.1) were immobilized onto the streptavidin sensor tip, and the experiments were performed using various concentrations of mature (A) and pro (B) MMP13 in HEPES buffered saline. Red lines represent fitted curve. Data are representative of at least three separate assays.

### HS induces dimerization of mature MMP13 but not proMMP13

To examine whether MMP13-HS interaction alters the oligomeric state of MMP13, we used size exclusion chromatography (SEC) combined with mass photometry. Incubating proMMP13 with either HS 10mer or 12mer did not induce any shift in the elution profile of proMMP13, suggesting these HS oligosaccharides cannot induce proMMP13 to form oligomers (Fig. 3A). In contrast, we found that binding of HS-10mer resulted in a slightly earlier elution position of mature MMP13 (14.9ml) compared to mature MMP13 alone (15.1ml); while binding of HS-12mer resulted in an even earlier elution position of mature MMP13 (Fig. 3B). This result suggests that HS 12mer likely induces dimerization of mature MMP13, while HS 10mer likely forms a stable complex with mature MMP13 without inducing dimerization. To further validate the dimerization of MMP13 induced by HS, we performed mass photometry and found the molecular weight of HS-12mer /mature MMP13 complex (86KDa ± 19.3KDa) is around 2-fold bigger than mature MMP13 (55KDa ± 10.2KDa), supporting the dimerization of mature MMP13 (Fig. 2C).

**Figure 3.**
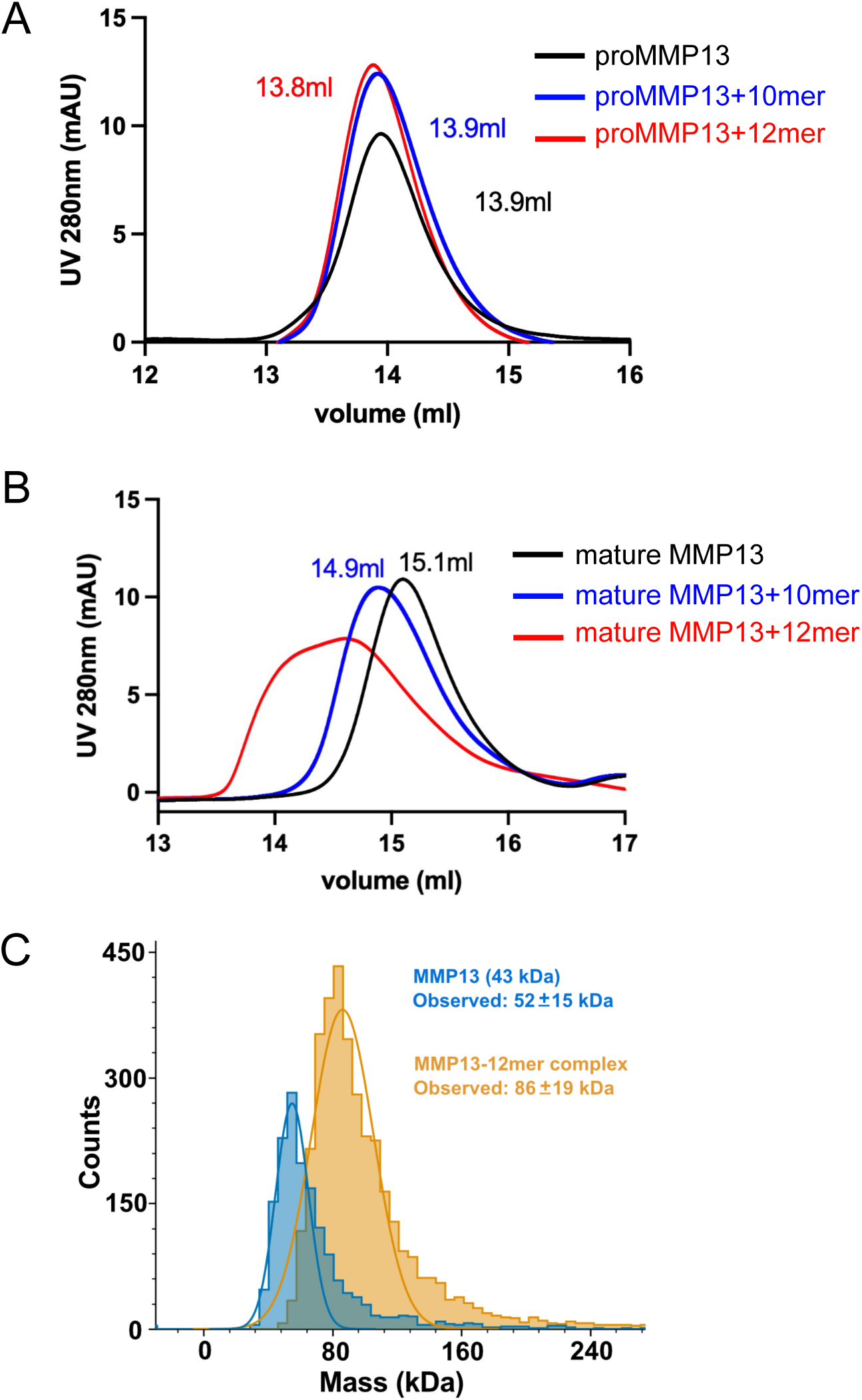
HS induces dimerization of mature MMP13 but not proMMP13. Size-exclusion chromatography of pro-MMP13 (A) and mature MMP13 (B) alone or after incubation with HS-10mer NS2S6S and-12mer NS2S6S on Superdex200 increase column. (C) Mass distributions of mature MMP13 monomer (blue line) and MMP13-12mer NS2S6S complex (yellow line) at 50 nM concentration. Peaks were assigned according to the predicted molecular mass of the MMP13 monomer and oligomeric species. Peak positions indicate the fitted molecular masses, and data are presented as the mean SD from three independent experiments.

### Binding of HS promotes the stability of mature MMP13

HS can regulate the stability of HS-binding proteins through two mechanisms: first, alteration of conformational stability; second, changing the susceptibility to proteolytic degradation. Using a peptidase assay, we found that incubating mature MMP13 at 37 °C resulted in a gradual loss of enzymatic activity, which suggests that prolonged incubation at physiological temperature is detrimental to the conformational stability of mature MMP13. Interestingly, in the presence of heparin, the peptidase activity of MMP13 was completely preserved (Fig. 4A, 4B), suggesting binding of heparin helps preserve the active conformation of MMP13.

**Figure 4.**
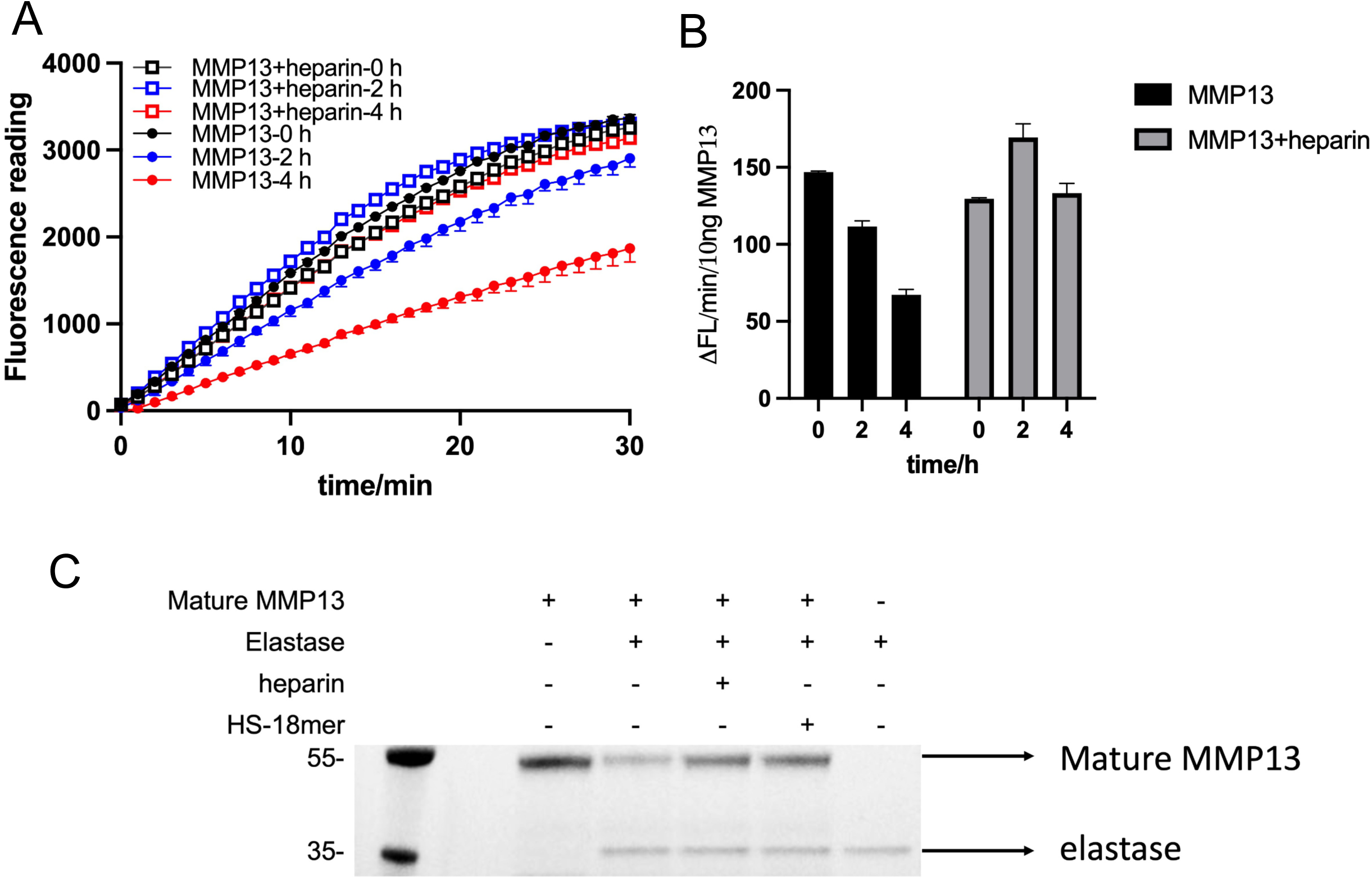
Binding of HS promotes the stability of mature MMP13. (A) Peptidase activity of mature MMP13 after incubation at 37 C in the presence or absence of heparin (1:2 molar ratio), measured using MCA-KPLGL-Dap(Dnp)-AR-NH_2_. (B) Initial activity rate calculated from the first 5 min of the progress curves in panel A. (C) Digestion of 5 µg mature MMP13 by 2 µg neutrophil elastase in the presence or absence of heparin or HS-18mer NS2S6S (1:10 molar ratio), visualized by silver staining

We further examined the effect of HS on proteolytic degradation of mature MMP13. To this end we digested mature MMP13 by elastase to mimic the in vivo degradation process with or without HS. We found heparin and HS-18mer NS2S6S significantly protected mature MMP13 from being digested by elastase, indicating that HS could prolong the half-life of mature MMP13 in vivo (Fig. 4C). Together, our data show that HS stabilizes the conformation of pro-MMP13 and protects mature MMP13 from degradation.

### Heparin has no effect on the activation of proMMP13

As HS is known to promote autoactivation of proteases such as MMP7 and cathepsin K, we investigated whether heparin has any effect on the activation process of MMP13^40,44,47^. Different from proMMP7 and proCathepsin K, we found that autoactivation mechanism is absent in activation of proMMP13 under normal conditions. Even when mature MMP13 is present in large access (5 molar excess), we observed no processing of proMMP13 (Fig. 5A). Activation of proMMP13 can only be achieved through proteolytic digestion by another protease, such as trypsin; or by compounds that disrupts the “cysteine switch”, such as 4-Aminophenylmercuric acetate (APMA), which allow autoactivation to proceed^48^.

**Figure 5.**
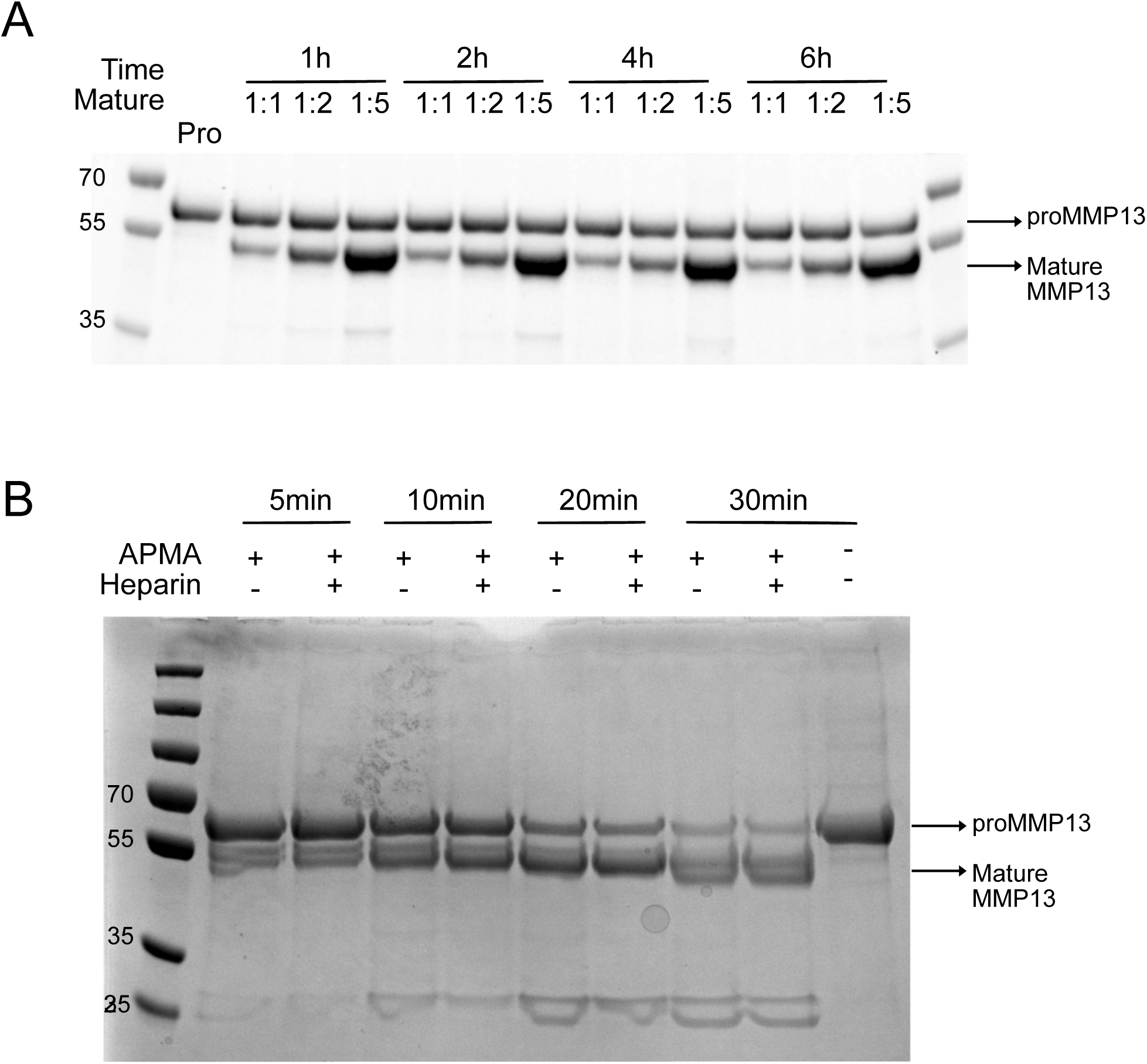
Heparin has no effect on the activation of proMMP13. (A) Processing of proMMP13 (10 µg) was performed by incubating with increasing amounts of mature MMP13 (10-50 µg) for 1 to 6 h. (B) ProMMP13 was activated by 1mM APMA in the presence or absence of heparin (1:1 molar ratio). Reactions were performed at 37 C for up to 30 min. The processing was monitored by Coomassie blue staining. Data are representative of at least three separate assays.

To determine whether HS regulates the activation process of MMP13, we incubated pro-MMP13 with 1mM APMA in the presence or absence of heparin (Fig. 5B). No significant difference was observed in the pro-MMP13 activation process with or without heparin. Similarly, presence of heparin did not alter the kinetics of trypsin-mediated activation of proMMP13 (Fig. 5C). Combined, our data showed that HS does not regulate the activation process of proMMP13.

### HS selectively inhibits the collagenase activity of mature MMP13

To determine whether HS binding affects the enzymatic activity of MMP13, we examined the proteolytic activity of MMP13 on digesting different substrates, including non-collagenous and collagen substrates. First, we tested a fluorescence-conjugated peptide substrate MCA-KPLGL-Dap(Dnp)-AR-NH_2_, and found heparin did not affect the peptide digestion by MMP13 (Fig. 6A and B). We also tested whether the presence of HS can affect digestion of non-collagen substrates by incubating mature MMP13 with osteoprotegrin and fibronectin, two known substrates of MMP13 (Fig. 6C). Again, heparin did not show any effect on digestion of these two substrates by mature MMP13. Together, our data suggests that HS does not directly affect the peptidase activity of MMP13.

**Figure 6.**
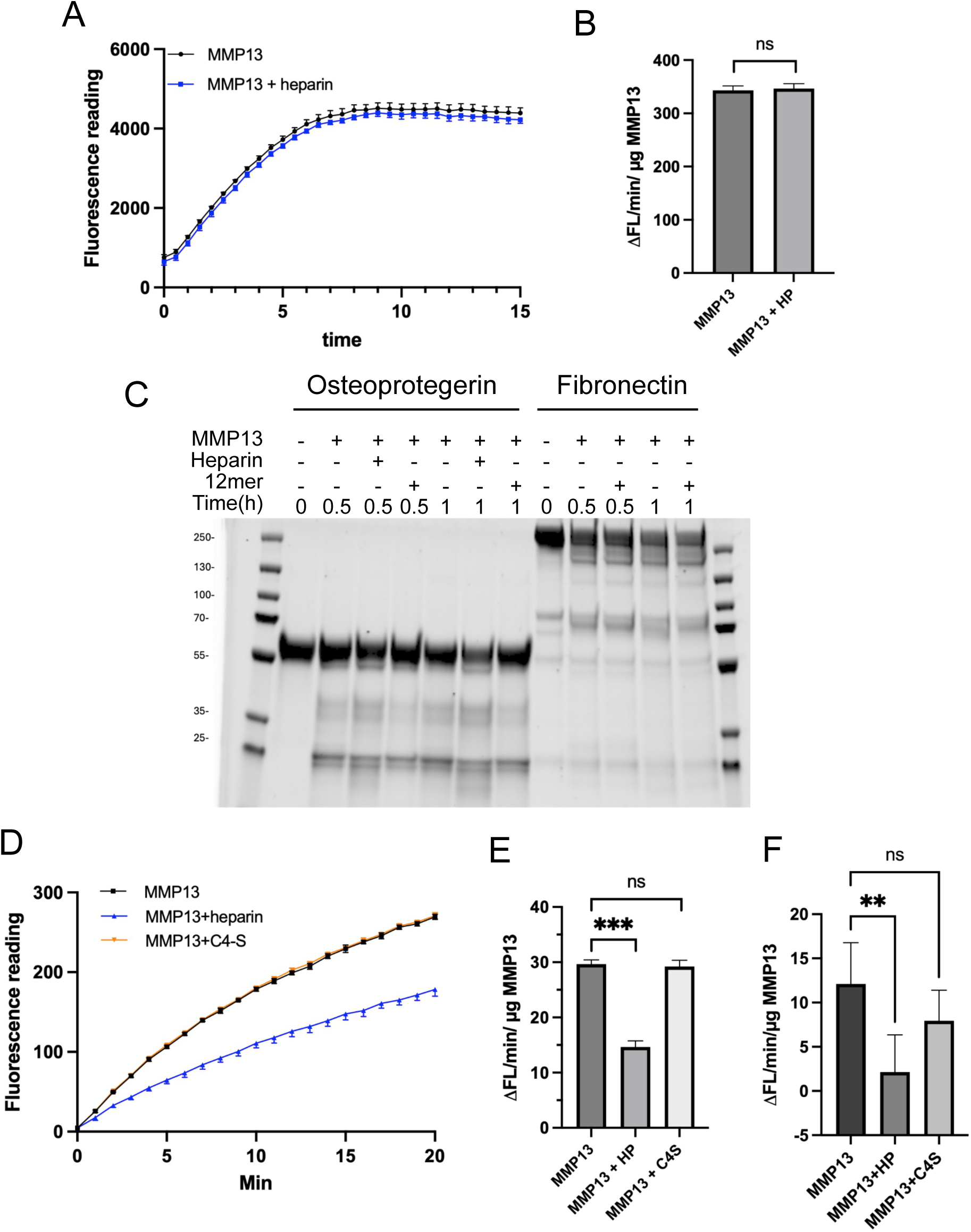
Heparin selectively inhibits the collagenase activity of mature MMP13. (A) Digestion of peptide substrate MCA-KPLGL-Dap (Dnp)-AR-NH2 with MMP13 (10 ng) in the presence or absence of heparin (30 ng). (B) Initial reaction rate calculated from the first 5 min of the progress curves in panel A. (C) Digestion of two extracellular proteins with MMP13 in the presence or absence of heparin or HS-12mer (compound #74). Left, 5 µg osteoprotegrin digestion by 1 µg MMP13 with heparin (1:10 molar ratio) or HS-12mer (1:5 molar ratio); Right, 5 µg fibronectin digestion by 50 ng MMP13 with heparin (1:10 molar ratio) or HS-12mer (1:5 molar ratio). (D) Digestion of FITC-labeled type I collagen with 500 ng MMP13 in the presence or absence of heparin and chondroitin A (C4-S) (1:1 molar ratio). (E) Initial reaction rate (first 5 min) of MMP13 derived from plot D. (F) Digestion of FITC-labeled type II collagen with 75 ng MMP13 in the presence or absence of heparin or C4-S (1:1 molar ratio). n = 3 technical replicates, *** represents p < 0.001, ** represents p < 0.01 by Student’s t test. Data are representative of at least three separate assays.

To examine whether the collagenase activity of MMP13 is affected by HS, we tested both FITC-labeled type I and type II collagen, two main collagen substrates for MMP13. We discovered for the first time that HS could inhibit the collagenase activity of MMP13 towards both type I (Fig. 6D and E) and II collagen (Fig. 6F). In contrast, chondroitin-4-sulfate, a highly abundant glycosaminoglycan found in the extracellular matrix, shows no inhibition of the collagenase activity of MMP13 (Fig. 6D and E), indicating that the inhibition is not pure electrostatic-driven. Together, these findings strongly suggest that HS could selectively inhibit the collagenase activity of MMP13.

Next, using a panel of structurally defined HS oligosaccharides, we examined the dependence of chain length and sulfation pattern on the inhibitory activity of HS. By comparing HS oligosaccharides of different lengths (10-, 12-, 14-, and 18-mers), we found that larger oligosaccharides inhibited MMP13 collagenase activity more effectively (Fig. 7A). Furthermore, we also tested HS 12mers with different sulfation patterns (compound #84 and #74) and found that one additional 3-O-sulfation could significantly increase its inhibitory potency on MMP13 collagenase activity (Fig. 7B). In addition, HS oligosaccharides inhibit the collagenase activity in a dose-dependent manner (Fig. 7C). We also examined the effect of HS on the type II collagen digestion by MMP13 and found HS could also significantly inhibit its digestion by MMP13, but the inhibition requires a 18mer (Fig. 7D).

**Figure 7.**
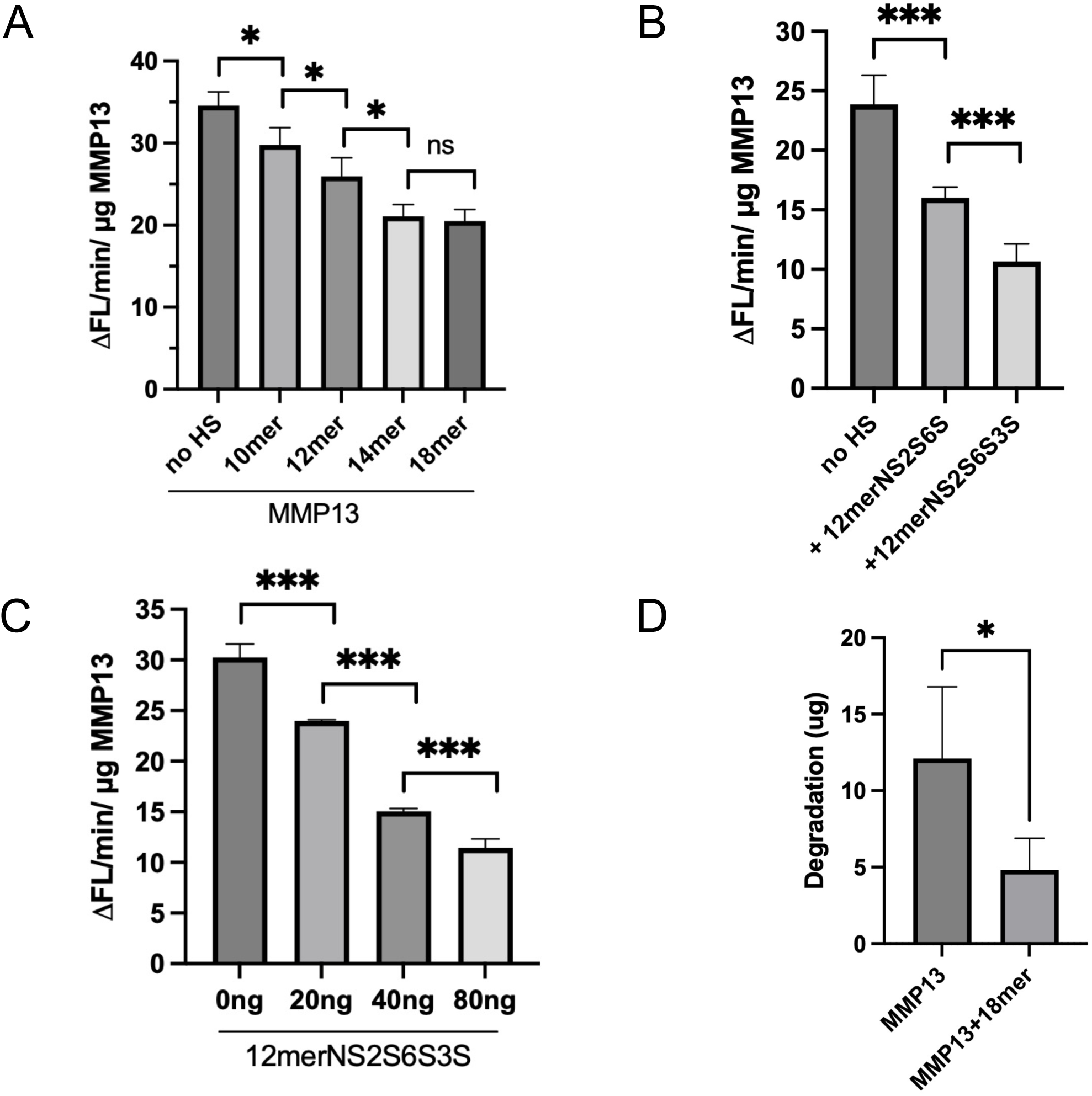
HS inhibits the collagenase activity of MMP13 in chain length-, sulfation-, and dose-dependent manners. (A) Digestion of FITC-labeled type I collagen with 500 ng MMP13 in the presence or absence of fully sulfated HS oligosaccharides with different chain sizes (10mer NS2S6S, 12mer NS2S6S, 14mer NS2S6S, and 18mer NS2S6S; at 1:1 molar ratio). (B) Digestion of FITC-labeled type I collagen with 500 ng MMP13 in the presence or absence of different HS-12mer (NS2S6S and NS2S3S6S; at 1:1 molar ratio). (C) Digestion of FITC-labeled type I collagen with 500 ng MMP13 in the presence or absence of HS-12mer NS2S3S6S (20-80 ng). (D) Digestion of FITC-labeled type II collagen with 75 ng MMP13 in the presence or absence of HS-18mer NS2S6S at a 1:1 molar ratio. n = 3 technical replicates, *** represents p < 0.001, ** represents p < 0.01, * represents p < 0.05 by Student’s t test. Data are representative of at least three separate assays.

### proMMP13 and mature MMP13 engage HS through partially distinct binding surfaces

The crystal structure of the mature MMP13 revealed a highly positively charged surface that spans both the catalytic and hemopexin domains, with contributions also coming from the linker sequence (Fig. 8A). This positively charged surface likely encompasses the HS binding site. To map the HS-binding site in MMP13, we performed site-directed mutagenesis of basic residues distributed across the catalytic domain, linker region, and hemopexin domain. The mutants were screened for binding to heparin-Sepharose column. According to the electrostatic surface potential of MMP13, we mutated conserved basic residues in the catalytic domain (Lys136, Lys139, Lys140, and Lys249), linker region (Lys276 and Lys279), and hemopexin domain (Lys304, Arg306, Lys324, Arg333, Arg350, Arg352, Lys353, Lys369, Lys377, Lys380, and Lys381).

**Figure 8.**
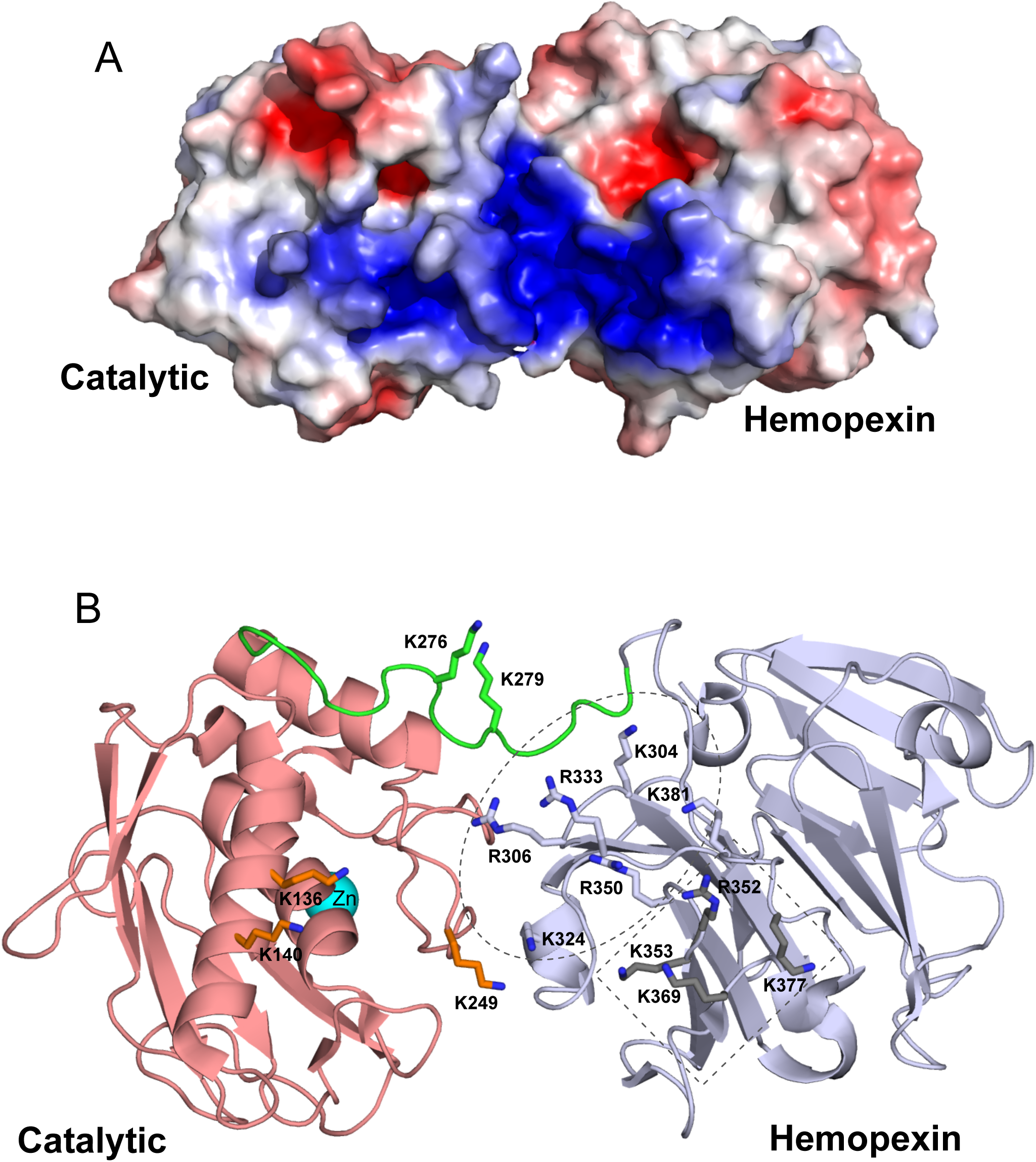
ProMMP13 and mature MMP13 engage HS through partially distinct binding surfaces. (A) Surface electrostatic potential of mature human MMP13 (PDB:4fvl) prepared in PyMOL. Positively charged surface is shown in blue and negatively charged surface is shown in red. (B) Side chains of residues potentially involved in binding to HS are shown in sticks. The active site of MMP13 is indicated by the catalytic Zn ion in cyan, which is located at the opposite surface of the current view. The two clusters of HS-binding residues in the hemopexin domain are marked in dashed oval (cluster-1) and dashed rectangle (cluster-2), respectively.

Analysis of the proenzyme mutants identified the hemopexin domain as the predominant HS-binding region (Table 1). Ten of the 11 hemopexin-domain mutants exhibited reduced NaCl concentration required for elution by at least 50 mM. The largest reductions were observed for K304A, R306A, R333A, R350Q, R352A, K353A, and K369A, which were eluted at NaCl concentrations 100–160 mM lower than WT pro-MMP13. By contrast, mutations in the catalytic domain had little effect on heparin binding. K276Q was the only linker-region mutation with reduced binding, displaying a reduction of the elution salt concentration by 75 mM. Combining R352A and K353A produced a larger reduction than both single mutations, decreasing the NaCl concentration required for elution by approximately 250 mM.

**Table 1.**
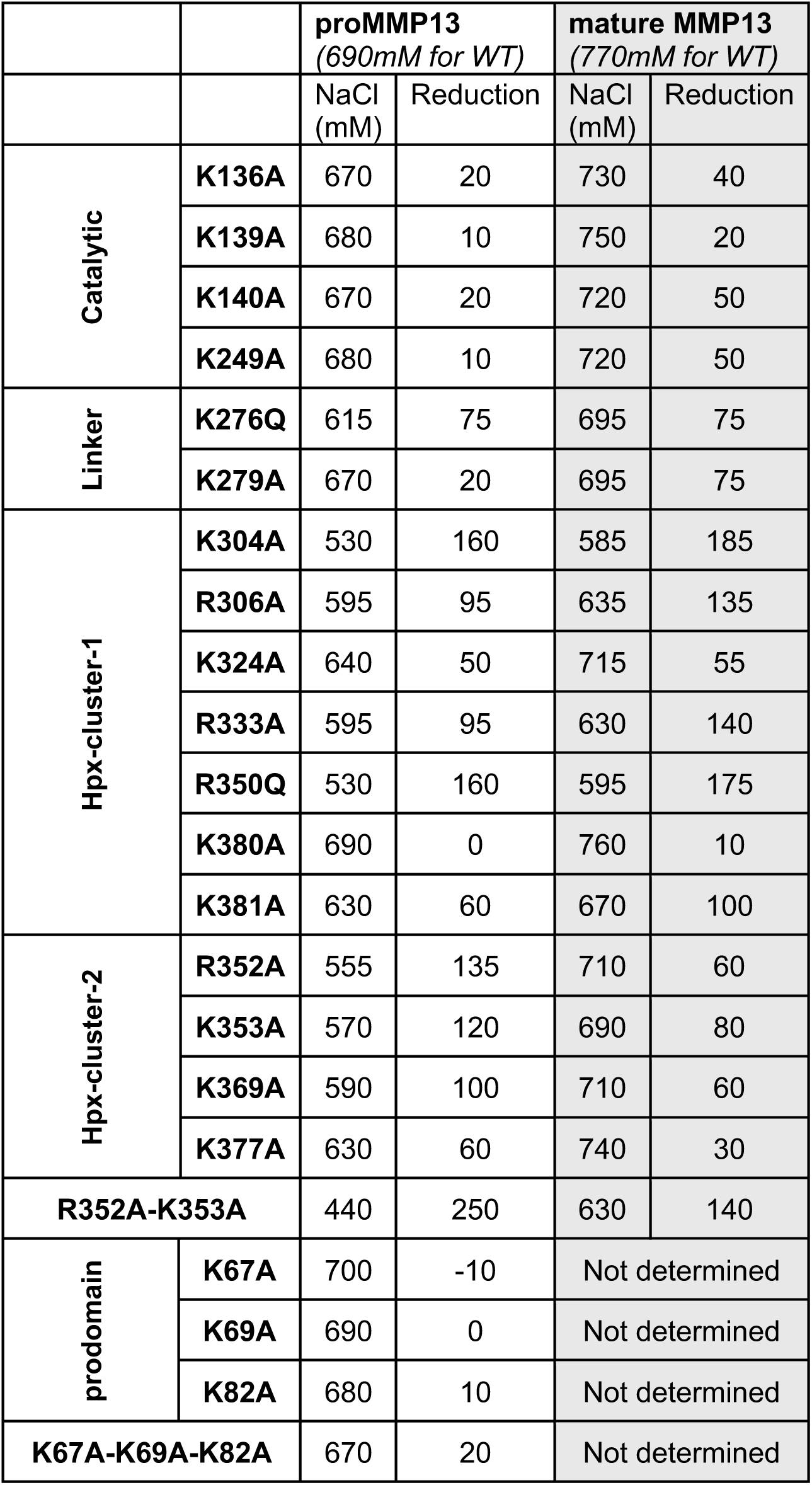
Concentration of salt required to elute MMP13 mutants from Heparin Sepharose column.

Next, we processed all proenzyme mutants into mature MMP13 and determined the impact of mutations on binding of mature MMP13 to heparin Sepharose column. Interestingly, while we found that the basic residues in the hemopexin domain still make the dominant contribution to the binding of mature enzyme, the relative importance of some of the residues changed significantly (Table 1). For the more centrally located cluster of basic residues (Hpx cluster-1, circled in dashed oval in Fig. 8B), including K304, R306, K324, R333, R350 and K381, their contribution to the mature MMP13-HS interaction is similar or slightly enhanced compared to their contribution to the proMMP13-HS interaction. In sharp contrast, the contribution to the mature MMP13-HS interaction was dramatically reduced for Hpx cluster-2 (marked in dashed rectangle in Fig. 8B, including R352, K353, K369 and K377) compared to their contribution to the proMMP13-HS interaction. In addition, we found several catalytic domain and linker region residues, including K136, K140, K249 and K279, which make only marginal contribution to the binding in proMMP13, are now making significant contribution to mature MMP13-HS interaction. These findings corroborate nicely with our finding on the different binding specificities of proMMP13 and mature MMP13 observed by our microarray analysis. Together, these data strongly suggest that HS likely interacts with proMMP13 and mature MMP13 in two very different modes with different involvement of binding residues.

Intrigued by our finding that the minimal length requirement of HS oligosaccharide for binding to proMMP13 is markedly longer than for binding to mature MMP13 (12mer versus 8mer, Fig. 1B, C), we tested the possibility that additional HS-binding residues might resides in the prodomain, which would effectively extend the length of the HS-binding site in proMMP13. We identified three conserved basic residues in the prodomain (Lys67, Lys69, and Lys82) that are in the same surface of the identified HS-binding site in the mature MMP13. However, neither the single mutations of these residues, nor the K67A-K69A-K82A triple mutants, had much impact on heparin binding (Table 1). This result indicates that the prodomain does not participate in proMMP13-HS interactions.

### HS-binding induces a significant conformational change of proMMP13

Using small-angle X-ray scattering (SAXS) technique, we examined whether binding of HS-oligosaccharide alters the conformation of proMMP13 by collecting data on proMMP13 alone and proMMP13-12mer complex. HS 12mer (compound 74) binding clearly induced a conformational change of proMMP13, resulting in a quite different scattering curve (Fig. 9A) and P(r) function plot (Fig. 9B). P(r) plot is used to describe the paired-set of distances between all the electrons within the protein, and the shape of which is directly determined by protein conformation. The P(r) plots show substantial differences between the two structures (Fig. 9B). From this plot we found that the R_g_ and D_max_ values of the proMMP13-12mer complex were both reduced ∼12% compared to proMMP13, suggesting that the complex adopt a more compact structure than proMMP13 in its apo form.

**Figure 9.**
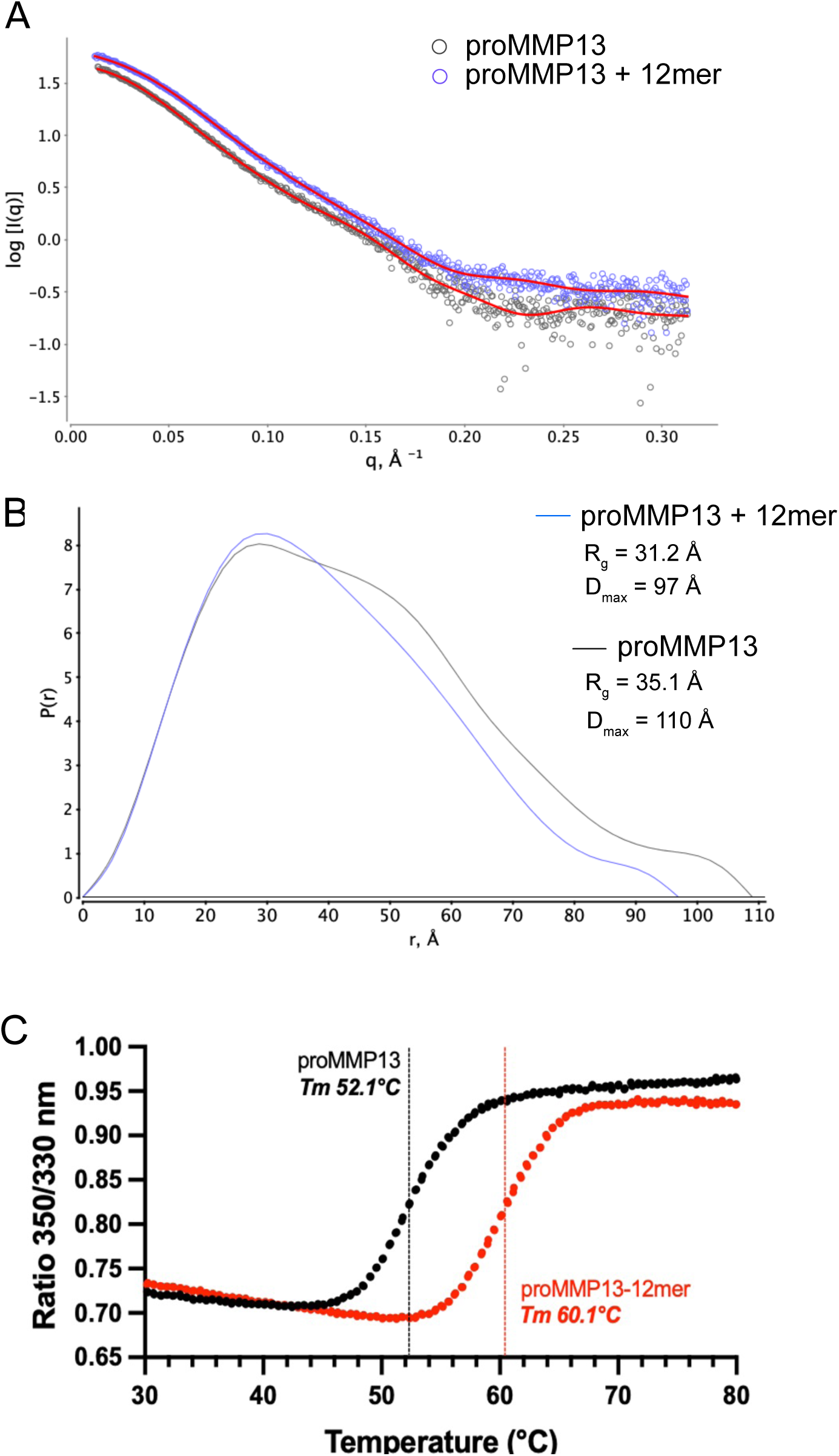
HS-binding induces a significant conformational change of proMMP13. Structural analysis of proMMP13/12mer complex by Small-angle X-ray scattering (SAXS). (A) Raw SAXS scattering curves proMMP13 or proMMP13-12mer complex. Red lines represent the fitted scattering curves. Data was collected at 5 mg/ml. B, P(r) function plots of proMMP13 and proMMP13-12mer complex. (C) NanoDSF thermograms of proMMP13 in the presence or absence of HS 12mer NS2S6S at a 1:1 molar ratio. Thermal unfolding was monitored using the intrinsic tryptophan fluorescence ratio at 350 nm/330 nm. Data representative of three independent experiments.

To support this finding, we determined the thermostability of proMMP13 using nanoscale Differential Scanning Fluorimetry (nanoDSF) in the presence or absence of HS 12mer. Remarkably, we found 12mer binding resulted in a ∼8 °C increase in Tm value, indicating greatly enhanced thermostability of pro-MMP13 when it complexed with HS (Fig. 3B). This finding is consistent with our finding with SAXS, both suggesting that HS-binding significantly alters the conformation of proMMP13.

### Ablation of HS-binding had no effect on the activity of MMP13 but rendered MMP13 insensitive to HS-mediated inhibition of collagenase activity

The R352A-K353A double mutant was selected to test whether direct HS binding is required for HS-mediated inhibition of MMP13 collagenase activity. Both the proenzyme and mature forms of this mutant exhibited near-background binding to CHO-K1 cells (Fig. 10A and B). The mutant retained peptidase and collagenase activities comparable to those of WT MMP13, indicating that the mutations did not substantially alter catalytic function. HS 12mer markedly inhibited the collagenase activity of WT MMP13 but had no significant effect on the R352A-K353A mutant (Fig. 10C and D). This finding suggests that direct HS–MMP13 binding is required for HS-mediated inhibition of MMP13 collagenase activity.

**Figure 10.**
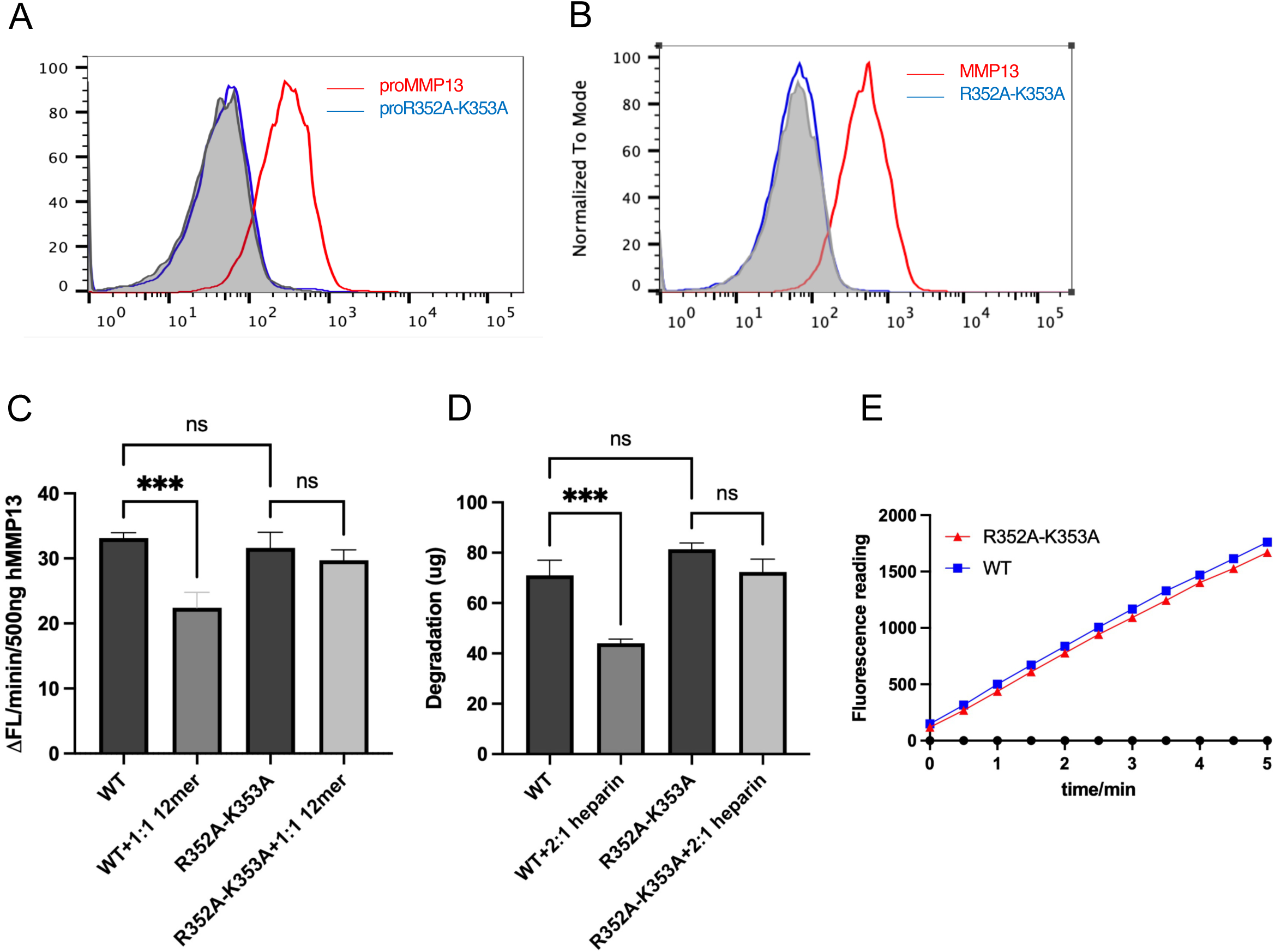
Ablation of HS-binding had no effect on the activity of MMP13 but renders MMP13 insensitive to HS-mediated inhibition of collagenase activity. Representative flow cytometry histograms of proMMP13 WT and R352A-K353A (A) or mature MMP13 (B) binding to CHO-K1 cells. The bound MMP13 was detected with a mouse anti-human MMP13 IgG, followed by anti-mouse-IgG Alexa-488. The shaded histogram is from cells stained with primary and secondary antibodies in the absence of MMP13. (C) Digestion of FITC-labeled type I collagen with 500 ng WT or mutant MMP13 in the presence or absence of HS-12mer NS2S3S6S (1:1 molar ratio). Plotted by the initial reaction rate (first 5 min). (D) Digestion of FITC-labeled type II collagen with 75 ng WT or mutant MMP13 in the presence or absence of heparin (2:1 molar ratio). (E) Digestion of peptide substrate MCA-KPLGL-Dap (Dnp)-AR-NH2 with WT and R352A-K353A MMP13 (10 ng). n = 3 technical replicates, *** represents p < 0.001 by Student’s t test. Data are representative of at least three separate assays.

## DISCUSSION

In this study, we aim to understand the molecular details of MMP13-HS interactions and the biochemical consequence of the interaction. We found that HS regulates the conformation, oligomerization state, stability and collagenase activity of MMP13. However, unlike MMP2 and MMP7, whose zymogen activation is regulated by HS^40–42^, zymogen activation of MMP13 is not affected by HS.

The HS-binding site of MMP13 is unique in that it involves more than 15 basic residues and spans an unusually large surface area of around 1200 Å^2^ (Fig. 8). This is much larger compared to the typical size of HS-binding sites, which are in the range of 300-500 Å^2^ and involve 5-10 basic residues^31^. Our mutagenesis study revealed that residues from the hemopexin domain, linker region and the catalytic domain all contribute to binding HS. However, the relative contribution of these HS-binding residues is different between proMMP13 and mature MMP13. We identified a core region, consisting of K304, R306, K324, R333, R350, K381 and K276, makes major contributions to binding of both proMMP13 and mature MMP13 to HS (Fig. 8B). The residues located in the catalytic domain (K136, K146, K249), and K279 in the linker region, play a more significant role in binding to mature MMP13 compared to proMMP13. In contrast, another cluster of basic residues located in the hemopexin domain (R352, K353, K369 and K377), which make major contributions in proMMP13-HS interaction, show sharply reduced contribution to mature MMP13-HS interaction. This data suggests that while the same set of residues are used in binding to proMMP13 and mature MMP13, they can adopt two distinct binding modes to HS depending on whether the prodomain is present. The two different binding modes resulted in markedly different preferences for HS structure (Fig. 1), and different binding kinetics of proMMP13 and mature MMP13 to HS (Fig. 2). Our study provided a novel insight on the plasticity of HS-binding sites. The biological implication of this alteration of binding characteristics of proMMP13 and mature MMP13 to HS remain unclear and is a major focus of our ongoing study.

Due to the triple helical nature of collagen, which is too bulky to fit into the active site of any protease, proteolytic digestion of collagen by collagenase requires intricate molecular mechanisms to facilitate digestion. Most collagenases recognize the triple-helical substrate with the help of at least one additional accessory domain, which helps locally unwind the triple helix and position an individual polypeptide chain within the catalytic cleft for hydrolysis^49–55^. In MMP this process requires cooperation between the catalytic and hemopexin domains. The most well-defined mechanism by which collagenases digest collagen fibers was presented in MMP-1^51,55^. This study suggests that for the bound collagen to unwind and fit into the active site of MMP1, the hemopexin and catalytic domains must flex with respect to each other to bend the bound collagen. Thus, the domain motion likely plays a critical role in facilitating structural changes of the collagen triple helix, which is required for collagenolysis. The structural feature that allows interdomain motion to occur is a linker peptide that connects the catalytic and hemopexin domains. Because our mutagenesis study has identified that the HS-binding site of MMP13 involves residues located in the hemopexin domain, catalytic domain and linker peptide, the most likely mechanism that contribute to the collagenase-specific inhibitory effect of HS is that binding of HS restricts the interdomain flexibility, thereby hampering the unwinding process that is required for collagenolysis. An alternative mechanism that might also contribute to the collagenase-specific inhibitory effect of HS is HS-induced dimerization of mature MMP13 (Fig. 3). Depending on the location of the dimerization interface, dimerization might interfere with the unwinding process of MMP13, or simply making the binding-unwinding cycle less efficient. The relative contribution of these two mechanisms remains a focal point of our current study.

It is now well established that MMP13 plays a significant role in the disease progression in OA. Clinical studies found that cartilage chondrocytes in OA patients express much higher level of MMP13 compared to chondrocytes in healthy joint^14,26^. Deletion of *Mmp13* protects against cartilage degeneration in experimental OA models^56^. In addition, postnatal overexpression of MMP13 in mice cartilage resulted in similar joint defects to those of human OA^57^. Given the essential role of MMP13 in cartilage degradation, it has been pursued as a promising therapeutic target in treating OA^58,59^. Early MMP inhibitors were designed to chelate the catalytic zinc ion. Because of the high structural similarity among MMPs, these inhibitors lacked selectivity and caused musculoskeletal syndrome^60^. Although MMP13-selective active site inhibitors were later developed, none has been successfully advanced to clinic^61–63^. While this lack of success can be attributed to multiple factors, such as poor pharmacokinetics and design of clinical trials, another possible factor could be our incomplete understanding the physiological role of MMP13 in cartilage homeostasis. In addition to its pathological role in collagen degradation, MMP13 might promote chondrocyte homeostasis by processing non-collagenous substrates. If that is the case, complete inhibition of the MMP13 activity might be undesirable and an ideal inhibitor might be the one that suppresses the collagenase activity of MMP13 while preserving its physiological activity toward other substrates.

The fact that binding of HS in between the hemopexin and catalytic domains inhibits the collagenase activity of MMP13 revealed a novel exosite of MMP13 that might be exploited to develop collagenase-specific inhibitors of MMP13. In principle, molecules that simultaneously bind the hemopexin and catalytic domains, the hemopexin domain and the linker peptide, or the catalytic domain and the linker peptide, have the potential to restrict the interdomain flexibility of MMP13 and inhibit MMP13 in a collagenase-specific manner. Due to the sheer distance between these structural elements, it’s unlikely that this can be achieved using small molecule inhibitors. In addition to HS oligosaccharides, we believe monoclonal antibodies (mAbs) should also be well suited to cover this molecular surface to restrict the interdomain flexibility MMP13. In theory, this can be achieved by developing mAbs that bind two domains simultaneously, or by designing bi-specific antibodies that bind the two domains separately.

In conclusion, our study identifies HS as an important regulator of the structure and function of MMP13. Future studies focusing on understanding the physiological importance of MMP13-HS interaction in vivo are expected to generate many new insights in MMP13 biology. Our data also strongly suggests that the HS-binding site of MMP13, which spans the catalytic domain, hemopexin domain and the linker region, represents a novel exosite that can be exploited to develop collagenase-specific inhibitors MMP13.

## MATERIEALS AND METHODS

### Expression and purification of full-length human proMMP13 in mammalian cells

Complete open reading frame of full-length human proMMP13 (GenScript) was cloned into the pcDNA3.1 (Thermo Fisher Scientific) expression vector using the XhoI and ApaI restriction sites. The final construct was verified by Sanger sequencing. Recombinant human pro-MMP13 was expressed by transient transfection of 293-Freestyle cells (ThermoFisher) by transient using FectoPRO transfection reagent (Polyplus) according to the manufacturer’s instructions. Conditioned medium was harvested 5 days after transfection, clarified by centrifugation at 5000 x g for 10 min, filtered through a 0.22 µm membrane, and purified by a HiTrap heparin-Sepharose column (Cytiva) equilibrated with HEPES buffer at pH 7.1. Bound protein was eluted using a linear NaCl gradient to 2 M. Purified pro-MMP13 was >99% pure as judged by silver staining.

### Activation of proMMP13 and purification of mature MMP13

proMMP13 activation was performed by using trypsin. Briefly, purified pro-MMP13 was diluted 2-fold with activation buffer containing 50mM Tris-HCl, 150mM NaCl, 5mM CaCl_2_, 0.01% Brij35, pH 7.5. Trypsin-EDTA (0.25%; Gibco) was added at a 1:200 volume ratio, and the mixture was incubated at 22 °C for 2 h. Activation was monitored using the fluorogenic substrate MCA-KPLGL-Dap(Dnp)-AR-NH_2_ (MedChemExpress) and by silver staining. Mature MMP13 was purified from the activation mixture by HiTrap heparin-Sepharose chromatography and eluted with a linear salt gradient from 150 mM to 2 M in MMP13 purification buffer containing 25 mM HEPES, 5 mM CaCl_2_, and 0.01%Brij35, pH7.5. Purified mature MMP13 was >99% pure as judged by silver staining.

### Heparin–Sepharose chromatography

To characterize the binding of proMMP13 and mature MMP13 to heparin, approximately 50 μg of purified protein was loaded onto a 1-mL HiTrap heparin–Sepharose column (Cytiva) equilibrated in MMP13 purification buffer. Bound protein was eluted with a linear salt gradient from 150 mM to 2 M at pH 7.5. The conductivity measurements at the elution peak were converted to NaCl concentration using a standard curve.

### Biolayer Interferometry Analysis

Interactions between HS and MMP13 were analyzed by biolayer interferometry using an Octet R8 instrument (Sartorius). Streptavidin biosensors (Sartorius) were hydrated and checked for 30 s before loading with 1 µg/ml biotinylated, fully sulfated HS-12mer (NS2S6S; compound#74) for 600 s. Immobilization of biotinylated glycosaminoglycans on streptavidin-coated sensor surfaces has been used previously for quantitative analysis of protein-glycosaminoglycan interactions^64^. All oligosaccharides and protein samples were prepared in assay buffer containing 25 mM HEPES, 150 mM NaCl, 5 mM CaCl_2_, and 0.01% Brij35, pH7.3. After HS immobilization, the sensors were equilibrated in buffer for 120 s to establish a baseline. Association was measured by transferring the HS-loaded sensors into solutions containing pro-MMP13 at 80, 100, 135, and 200 nM or mature MMP13 at 100, 200, 400, or 800 nM for 300 s. Dissociation was subsequently monitored in protein-free buffer for 300 s. Sensors were regenerated for 30 s between measurement cycles. Signals from blank sensors incubated in assay buffer with the same concentration range of proteins were subtracted from the corresponding binding responses. All measurements were performed at 30 °C with orbital shaking at 1000 rpm. Reference-corrected sensorgrams were aligned to the start of the association step and analyzed using the Octet Data Analysis software. Sensorgrams were fitted globally using a 1:1 Langmuir binding model with shared association and dissociation rate constants. The equilibrium dissociation constant was calculated as *K_D_* = *k*_off_/*k*_on_. Because HS promotes the dimerization of mature MMP13, kinetic parameters obtained from the 1:1 fit for mature MMP13 were interpreted as apparent values rather than as parameters of a strictly bimolecular interaction. Fit quality was assessed from the coefficient of determination, reduced chi-square value, and residual distribution. Measurements were performed in three independent experiments, with 3 technical replicates and kinetic parameters were reported as the mean ± SD.

### Alexa Fluor 488 labeling of pro-MMP13 and mature MMP13

Recombinant pro-MMP13 and mature MMP13 were labeled with Alexa Fluor 488 NHS ester (Invitrogen) while bound to 200 µl packed heparin gravity columns. Pro-MMP13 (300 µg) was bound in 25 mM HEPES and 150 mM NaCl, pH 7.1, whereas mature MMP13 (250 µg) was bound in 25 mM HEPES, 150 mM NaCl, 5 mM CaCl₂, and 0.01% Brij35, pH 7.5. Column-bound proteins were incubated at 4 °C with 200 µL of Alexa Fluor 488 NHS ester at 0.5 mg/mL for 2 h. Reactions were quenched with 50 mM Tris-HCl, pH 7.7. Labeled proteins were eluted with high-salt buffer, diluted to approximately 400 mM NaCl, and further purified by HiTrap heparin-sepharose chromatography. Fractions eluting at the same positions as the corresponding unlabeled proteins were combined and concentrated.

### Glycan array binding assay

Amine-functionalized HS oligosaccharides were covalently immobilized through their reducing ends on NHS-functionalized glass slides, with each compound represented by 36 spots in a 6 × 6 pattern. The original array included HS structures varying in chain length and N-, 2-O-, 6-O-, and 3-O-sulfation. The microarray was probed with 1 µg/ml Alexa Fluor 488-labeled proMMP13 in 25 mM HEPES buffer containing 150 mM NaCl, pH7.3 or mature MMP13 in 25 mM HEPES containing 0.01%Brij35, 5 mM CaCl_2_, 150 mM NaCl, pH 7.5. After incubation and washing, the slides were scanned with a GenePix 4300A microarray scanner using a 488 nm laser. Fluorescence intensities were quantified using GenePix software. Binding to each HS oligosaccharide was reported as the mean fluorescence intensity ± SD of 36 spots per oligosaccharide.

### Analytical size-exclusion chromatography (SEC)

To determine the MMP13-HS complexes formation, purified pro-MMP13 or mature MMP13 (50 μg) was incubated with the indicated HS oligosaccharides at a 1:1 molar ratio overnight at 4 °C in 25 mM HEPES, 150mM NaCl, 5 mM CaCl_2,_ 0.01%Brij35, pH 7.5. Complexes were resolved at 4 °C on a Superdex 200 Increase column (Cytiva) equilibrated in the same buffer.

### Mass photometry

Mass photometry was performed using a Refeyn mass photometer to assess the oligomeric states mature MMP13 in the presence or absence of fully sulfated HS-12mer NS2S6S. Before sample acquisition, the instrument was calibrated using a four-protein molecular-mass standard diluted 500-fold in the same buffer as the corresponding MMP13 sample. Mature MMP13 was analyzed in 25 mM HEPES, 150 mM NaCl, 5 mM CaCl_₂_, and 0.01% Brij35, pH7.5. MMP13 was prepared at 1 µM with or without HS-12mer at a 1:1 molar ratio and diluted 20-fold before measurement. For droplet dilution, 10 µl of buffer was added to the slide well and focused before addition of 10 µl of diluted sample. Molecular-mass distributions were recorded and analyzed using Refeyn software.

### MMP13 stability assay

Mature MMP13 was stored alone or with heparin at a 1:5 molar ratio in 50 mM Tris-HCl, 150 mM NaCl, 5 mM CaCl2, and 0.01%Brij35, pH7.5, at 37 °C in for 4 h. The heparin used in this study was unfractionated porcine mucosal heparin (Scientific Protein Laboratories) with an average molecular mass of 15 kDa. 10 ng aliquots were taken per day or hour to measure the peptidase activity using the fluorogenic substrate as described above.

### MMP13 degradation assay

Mature MMP13 (5 µg) was incubated with elastase (2 µg, porcine pancreas, ThermoScientific) in activation buffer containing 50 mM Tris-HCl, 150 mM NaCl, 5 mM CaCl₂, and 0.01% Brij35, pH 7.5, at 22°C for 7 h. Reactions were performed in the absence or presence of heparin or fully sulfated HS-18mer NS2S6S at a 1:10 MMP13-to-glycosaminoglycan molar ratio. Reactions were terminated by addition of SDS-PAGE sample buffer and heat denaturation. Degradation products were separated by SDS-PAGE and visualized by silver staining.

### Small-angle X-ray scattering analysis

Scattering data will be collected at the SIBLYLS beamline at the Lawrence Berkeley National Laboratory through a mail-in SAXS service. ProMMP13-12mer complex was prepared by mixing 400 µg of SEC purified proMMP13 with 50 µg 12mer (molar ratio 1:2) overnight in HEPES buffer (25 mM HEPES, pH 7.1, 0.15 M NaCl, pH 7.1) at 4°C. The complex, and free proMMP13, were concentrated to 1 mg, 2 mg and 5 mg/mL for data collection at the SIBLYLS beamline 12.3.1. The data will be analyzed by SAXS analysis software Scatter 3. The whole set of experiments were performed twice using two different proMMP13 preparations with similar results.

### Nano Differential Scanning Fluorimetry

Nano-differential scanning fluorimetry (nanoDSF) was performed using a Prometheus Panta instrument (NanoTemper Technologies) to assess the thermal stability of pro-MMP13 and mature MMP13 in the presence or absence of fully sulfated HS-12mer NS2S6S. Pro-MMP13 was prepared at 500 µg/ml in 25 mM HEPES, 150 mM NaCl, pH 7.3. Mature MMP13 was prepared at 500 µg/ml in 25 mM HEPES, 150 mM NaCl, 5 mM CaCl₂, and 0.01% Brij35, pH 7.5. Where indicated, HS-12mer was added before sample loading. Samples (10 µl) were loaded into standard nanoDSF capillaries and heated from 25 °C to 80 °C at 1 °C/min. Tryptophan fluorescence emission was recorded at 330 and 350 nm after excitation at 280 nm. The 350/330 nm fluorescence ratio and its first derivative were calculated using ThermControl software.

### MMP13 peptidase activity assay

MMP13 peptidase activity was measured using the fluorogenic substrate MCA-KPLGL-Dap(Dnp)-AR-NH₂. Pro-MMP13 was activated by mixing 20 µl of protein with an equal volume of assay buffer containing 50 mM Tris-HCl, 150 mM NaCl, 5 mM CaCl₂, and 0.01% Brij35, pH 7.5. Trypsin-EDTA (0.25%) was added at a 1:200 dilution and samples were incubated at room temperature for 2 h. Activated MMP13 was diluted to 0.2 µg/ml in assay buffer. Each reaction contained 50 µl of mature enzyme and was initiated by adding 50 µl of substrate at 20 µM, yielding final concentrations of 0.1 µg/ml MMP13 and 10 µM substrate. Substrate-only wells served as blanks. Fluorescence was monitored in black 96-well plates using a SpectraMax plate reader (Molecular Devices) at 320 nm excitation and 405 nm emission for 30 min at 30 s intervals. Enzyme activity was calculated from the linear portion of each progress curve and normalized to the amount of MMP13.

Proteolysis of fibronectin and osteoprotegerin (human OPG) by mature MMP13 was assessed in vitro. Goat fibronectin (5 µg) was incubated with 50 ng mature MMP13 at 37 °C for 30 or 60 min in 50 mM Tris-HCl, 150 mM NaCl, 5 mM CaCl₂, and 0.01% Brij35, pH 7.5, with or without heparin at a 1:10 molar ratio or HS-12mer NS2S3S6S at a 1:5 molar ratio. Human OPG (5 µg) was incubated with 1 µg of mature MMP13 in the same buffer. Reactions were terminated with SDS-PAGE sample buffer and heat denaturation, and degradation was analyzed by SDS-PAGE.

### MMP13 collagenase activity assay

MMP13 collagenase activity was measured using FITC–labeled bovine type I collagen (ThermoFisher), FITC-labeled bovine type II collagen (Chondrex), and insoluble bovine type I collagen (Chondrex). For FITC-labeled type I collagen, mature MMP13 (500 ng) was combined with 5 µg of collagen in a final volume of 200 µl and incubated at 37 °C. Fluorescence was monitored for 1 h at 485 nm excitation and 525 nm emission, and activity was measured using a SpectraMax plate reader. Activity was calculated from the linear portion of the progress curve.

For FITC-labeled type II collagen, mature MMP13 was incubated with 100 µl of 1× FITC-labeled bovine type II collagen in activation buffer at 38 °C for 30 min. Reactions were terminated with 5 µl of 10 mM 1,10-phenanthroline, followed by addition of 5 µl of elastase and incubation at 38 °C for 20 min. Dioxane extraction buffer (200 µl) was added, samples were mixed and centrifuged at 10,000 rpm for 10 min, and 200 µl of each supernatant was transferred to a black 96-well plate. Fluorescence was measured at 490 nm excitation and 520 nm emission. For insoluble type I collagen, 2 mg of collagen was loaded into a 0.2 µm Nanosep centrifugal device, equilibrated for 15 min with 500 µl of buffer containing 50 mM Tris-HCl, 400 mM NaCl, 10 mM CaCl₂, 10 µM ZnCl₂, and 0.01% Brij35, pH 7.5 and centrifuged at 13,000 × g for 2 min. Collagen was incubated with 200 µl of 1 µM mature MMP13 for 2 h at room temperature. Filtrates were collected by centrifugation, and reactions were terminated with 38 mM EDTA. Collagen hydrolysis was quantified by fluorescamine derivatization and fluorescence measurement at 390 nm excitation and 475 nm emission.

The above reactions were carried out in the presence or absence of heparin, heparin-derived oligosaccharides, chondroitin sulfate A, or the indicated HS oligosaccharides at the lengths, sulfation patterns, and molar ratios specified in the corresponding figure legends.

### Site-directed mutagenesis

MMP13 mutants were prepared by site-directed mutagenesis, and the mutations were verified by Sanger sequencing. Expression, purification, and activation of the mutants were carried out the same way as the WT MMP13 described above. All mutants were expressed at a comparable level to WT MMP13 and showed comparable enzymatic activity to WT MMP13.

### Statistical analysis

All data are reported as means ± SDs. Statistical analysis was performed using GraphPad Prism software (GraphPad Software Inc.). The significance was assessed using two-tailed student’s t-tests or analysis of variance (ANOVA). P value< 0.05 was considered statistically significant.

## ACKNOWLEDGMENTS

SAXS data was collected at the Advanced Light Source (ALS) at the SIBYLS beamline, a national user facility operated by Lawrence Berkeley National Laboratory on behalf of the Department of Energy, Office of Basic Energy Sciences, through the Integrated Diffraction Analysis Technologies (IDAT) program, supported by DOE Office of Biological and Environmental Research. Additional support comes from the National Institute of Health project ALS-ENABLE (P30 GM124169). We also thank Maros Huliciak for his generous help with the mass photometry experiments. This work is supported by National Institutes of Health grant R01DE031273.

## CONFLICT OF INTEREST

J.L. is the founder of Glycan Therapeutics and has equity. GS is an employee of Glycan Therapeutics and has an equity option from the company. Other authors declare no competing interests.

## REFERENCE

1. Cui N, Hu M, Khalil RA. Biochemical and Biological Attributes of Matrix Metalloproteinases. In: Progress in Molecular Biology and Translational Science. Vol 147. Elsevier; 2017:1–73. doi:10.1016/bs.pmbts.2017.02.005

2. Sternlicht MD, Werb Z. How Matrix Metalloproteinases Regulate Cell Behavior. Annu Rev Cell Dev Biol. 2001;17(1):463–516. doi:10.1146/annurev.cellbio.17.1.463

3. De Almeida LGN, Thode H, Eslambolchi Y, et al. Matrix Metalloproteinases: From Molecular Mechanisms to Physiology, Pathophysiology, and Pharmacology. Pharmacol Rev. 2022;74(3):714–770. doi:10.1124/pharmrev.121.000349

4. Page-McCaw A, Ewald AJ, Werb Z. Matrix metalloproteinases and the regulation of tissue remodelling. Nat Rev Mol Cell Biol. 2007;8(3):221–233. doi:10.1038/nrm2125

5. Hey S, Linder S. Matrix metalloproteinases at a glance. J Cell Sci. 2024;137(2):jcs261898. doi:10.1242/jcs.261898

6. Chen K, Xu M, Lu F, He Y. Development of Matrix Metalloproteinases-Mediated Extracellular Matrix Remodeling in Regenerative Medicine: A Mini Review. Tissue Eng Regen Med. 2023;20(5):661–670. doi:10.1007/s13770-023-00536-x

7. Legendre C, Dayan G, Rousselle P. When proteases reshape barriers: Basement membrane remodelling in development, wound healing and tumour progression. FEBS J. 2026;293(13):3954–3972. doi:10.1111/febs.70585

8. Töpfer U, Dahlitz I, Holz A. Mmp2 regulates basement membrane remodeling and dedieerentiation of the visceral musculature during Drosophila metamorphosis. Sci Rep. 2026;16(1):7827. doi:10.1038/s41598-026-41763-1

9. Gkouveris I, Nikitakis N, Aseervatham J, Rao N, Ogbureke K. Matrix metalloproteinases in head and neck cancer: current perspectives. Met Med. 2017;Volume 4:47–61. doi:10.2147/MNM.S105770

10. Löeek S, Schilling O, Franzke CW. Biological role of matrix metalloproteinases: a critical balance. Eur Respir J. 2011;38(1):191–208. doi:10.1183/09031936.00146510

11. Fanjul-Fernández M, Folgueras AR, Cabrera S, López-Otín C. Matrix metalloproteinases: Evolution, gene regulation and functional analysis in mouse models. Biochim Biophys Acta BBA - Mol Cell Res. 2010;1803(1):3–19. doi:10.1016/j.bbamcr.2009.07.004

12. Madzharova E, Kastl P, Sabino F, Auf Dem Keller U. Post-Translational Modification-Dependent Activity of Matrix Metalloproteinases. Int J Mol Sci. 2019;20(12):3077. doi:10.3390/ijms20123077

13. Rose BJ, Kooyman DL. A Tale of Two Joints: The Role of Matrix Metalloproteases in Cartilage Biology. Dis Markers. 2016;2016:1–7. doi:10.1155/2016/4895050

14. Billinghurst RC, Dahlberg L, Ionescu M, et al. Enhanced cleavage of type II collagen by collagenases in osteoarthritic articular cartilage. J Clin Invest. 1997;99(7):1534–1545. doi:10.1172/JCI119316

15. Knäuper V, López-Otin C, Smith B, Knight G, Murphy G. Biochemical Characterization of Human Collagenase-3. J Biol Chem. 1996;271(3):1544–1550. doi:10.1074/jbc.271.3.1544

16. Mitchell PG, Magna HA, Reeves LM, et al. Cloning, expression, and type II collagenolytic activity of matrix metalloproteinase-13 from human osteoarthritic cartilage. J Clin Invest. 1996;97(3):761–768. doi:10.1172/JCI118475

17. Zhang C, Tang W, Li Y. Matrix Metalloproteinase 13 (MMP13) Is a Direct Target of Osteoblast-Specific Transcription Factor Osterix (Osx) in Osteoblasts. Wu Q, ed. PLoS ONE. 2012;7(11):e50525. doi:10.1371/journal.pone.0050525

18. Lausch E, Keppler R, Hilbert K, et al. Mutations in MMP9 and MMP13 Determine the Mode of Inheritance and the Clinical Spectrum of Metaphyseal Anadysplasia. Am J Hum Genet. 2009;85(2):168–178. doi:10.1016/j.ajhg.2009.06.014

19. Kennedy AM, Inada M, Krane SM, et al. MMP13 mutation causes spondyloepimetaphyseal dysplasia, Missouri type (SEMDMO). J Clin Invest. 2005;115(10):2832–2842. doi:10.1172/JCI22900

20. Takaishi H, Kimura T, Dalal S, Okada Y, D’Armiento J. Joint Diseases and Matrix Metalloproteinases: A Role for MMP-13. Curr Pharm Biotechnol. 2008;9(1):47–54. doi:10.2174/138920108783497659

21. Inada M, Wang Y, Byrne MH, et al. Critical roles for collagenase-3 (Mmp13) in development of growth plate cartilage and in endochondral ossification. Proc Natl Acad Sci. 2004;101(49):17192–17197. doi:10.1073/pnas.0407788101

22. Stickens D, Behonick DJ, Ortega N, et al. Altered endochondral bone development in matrix metalloproteinase 13-deficient mice. Development. 2004;131(23):5883–5895. doi:10.1242/dev.01461

23. Johansson N, Saarialho-Kere U, Airola K, et al. Collagenase-3 (MMP-13) is expressed by hypertrophic chondrocytes, periosteal cells, and osteoblasts during human fetal bone development. Dev Dyn. 1997;208(3):387–397. doi:10.1002/(SICI)1097-0177(199703)208:3<387::AID-AJA9>3.0.CO;2-E

24. Yamamoto K, Okano H, Miyagawa W, et al. MMP-13 is constitutively produced in human chondrocytes and co-endocytosed with ADAMTS-5 and TIMP-3 by the endocytic receptor LRP1. Matrix Biol. 2016;56:57–73. doi:10.1016/j.matbio.2016.03.007

25. Wang X, Manner PA, Horner A, Shum L, Tuan RS, Nuckolls GH. Regulation of MMP-13 expression by RUNX2 and FGF2 in osteoarthritic cartilage. Osteoarthritis Cartilage. 2004;12(12):963–973. doi:10.1016/j.joca.2004.08.008

26. Vincenti MP, Brinckerhoe CE. Transcriptional regulation of collagenase (MMP-1, MMP-13) genes in arthritis: integration of complex signaling pathways for the recruitment of gene-specific transcription factors. Arthritis Res Ther. 2002;4(3):157. doi:10.1186/ar401

27. Li H, Wang D, Yuan Y, Min J. New insights on the MMP-13 regulatory network in the pathogenesis of early osteoarthritis. Arthritis Res Ther. 2017;19(1):248. doi:10.1186/s13075-017-1454-2

28. Aida Y, Maeno M, Suzuki N, Shiratsuchi H, Motohashi M, Matsumura H. The eeect of IL-1β on the expression of matrix metalloproteinases and tissue inhibitors of matrix metalloproteinases in human chondrocytes. Life Sci. 2005;77(25):3210–3221. doi:10.1016/j.lfs.2005.05.052

29. Akkiraju H, Nohe A. Role of Chondrocytes in Cartilage Formation, Progression of Osteoarthritis and Cartilage Regeneration. J Dev Biol. 2015;3(4):177–192. doi:10.3390/jdb3040177

30. Karamanos NK, Theocharis AD, Piperigkou Z, et al. A guide to the composition and functions of the extracellular matrix. FEBS J. 2021;288(24):6850–6912. doi:10.1111/febs.15776

31. Xu D, Esko JD. Demystifying Heparan Sulfate–Protein Interactions. Annu Rev Biochem. 2014;83(1):129–157. doi:10.1146/annurev-biochem-060713-035314

32. Severmann AC, Jochmann K, Feller K, et al. An altered heparan sulfate structure in the articular cartilage protects against osteoarthritis. Osteoarthritis Cartilage. 2020;28(7):977–987. doi:10.1016/j.joca.2020.04.002

33. Gerstner M, Severmann AC, Chasan S, Vortkamp A, Richter W. Heparan Sulfate Deficiency in Cartilage: Enhanced BMP-Sensitivity, Proteoglycan Production and an Anti-Apoptotic Expression Signature after Loading. Int J Mol Sci. 2021;22(7):3726. doi:10.3390/ijms22073726

34. Shamdani S, Chantepie S, Flageollet C, et al. Heparan sulfate functions are altered in the osteoarthritic cartilage. Arthritis Res Ther. 2020;22(1):283. doi:10.1186/s13075-020-02352-3

35. Chanalaris A, Clarke H, Guimond SE, Vincent TL, Turnbull JE, Troeberg L. Heparan Sulfate Proteoglycan Synthesis Is Dysregulated in Human Osteoarthritic Cartilage. Am J Pathol. 2019;189(3):632–647. doi:10.1016/j.ajpath.2018.11.011

36. Koyama Y, Naruo H, Yoshitomi Y, et al. Matrix Metalloproteinase-9 Associated with Heparan Sulphate Chains of GPI-Anchored Cell Surface Proteoglycans Mediates Motility of Murine Colon Adenocarcinoma Cells. J Biochem (Tokyo). 2008;143(5):581–592. doi:10.1093/jb/mvn006

37. Ruiz-Gómez G, Vogel S, Möller S, Pisabarro MT, Hempel U. Glycosaminoglycans influence enzyme activity of MMP2 and MMP2/TIMP3 complex formation - Insights at cellular and molecular level. Sci Rep. 2019;9(1):4905. doi:10.1038/s41598-019-41355-2

38. Tocchi A, Parks WC. Functional interactions between matrix metalloproteinases and glycosaminoglycans. FEBS J. 2013;280(10):2332–2341. doi:10.1111/febs.12198

39. Yu WH, Woessner JF. Heparan Sulfate Proteoglycans as Extracellular Docking Molecules for Matrilysin (Matrix Metalloproteinase 7). J Biol Chem. 2000;275(6):4183–4191. doi:10.1074/jbc.275.6.4183

40. Ra HJ, Harju-Baker S, Zhang F, Linhardt RJ, Wilson CL, Parks WC. Control of Promatrilysin (MMP7) Activation and Substrate-specific Activity by Sulfated Glycosaminoglycans. J Biol Chem. 2009;284(41):27924–27932. doi:10.1074/jbc.M109.035147

41. Crabbe T, Ioannou C, Docherty AJP. Human progelatinase A can be activated by autolysis at a rate that is concentration-dependent and enhanced by heparin bound to the C-terminal domain. Eur J Biochem. 1993;218(2):431–438. doi:10.1111/j.1432-1033.1993.tb18393.x

42. Koo BH, Han JH, Yeom YI, Kim DS. Thrombin-dependent MMP-2 Activity Is Regulated by Heparan Sulfate. J Biol Chem. 2010;285(53):41270–41279. doi:10.1074/jbc.M110.171595

43. Ryu HY, Lee J, Yang S, et al. Syndecan-2 Functions as a Docking Receptor for Pro-matrix Metalloproteinase-7 in Human Colon Cancer Cells. J Biol Chem. 2009;284(51):35692–35701. doi:10.1074/jbc.M109.054254

44. Fulcher YG, Prior SH, Masuko S, et al. Glycan Activation of a Sheddase: Electrostatic Recognition between Heparin and proMMP-7. Structure. 2017;25(7):1100–1110.e5. doi:10.1016/j.str.2017.05.019

45. Yu W hsuan, Woessner JF. Heparin-Enhanced Zymographic Detection of Matrilysin and Collagenases. Anal Biochem. 2001;293(1):38–42. doi:10.1006/abio.2001.5099

46. Murphy G, Nagase H. Localizing matrix metalloproteinase activities in the pericellular environment. FEBS J. 2011;278(1):2–15. doi:10.1111/j.1742-4658.2010.07918.x

47. Hao H, Zhang X, Liu J, Xu D. Heparan sulfate promotes autoactivation of procathepsin K by destabilizing the propeptide–catalytic domain interaction. Biochem J. 2026;483(6):967–980. doi:10.1042/BCJ20260109

48. Van Wart HE, Birkedal-Hansen H. The cysteine switch: a principle of regulation of metalloproteinase activity with potential applicability to the entire matrix metalloproteinase gene family. Proc Natl Acad Sci. 1990;87(14):5578–5582. doi:10.1073/pnas.87.14.5578

49. Stura EA, Visse R, Cuniasse P, Dive V, Nagase H. Crystal structure of full-length human collagenase 3 (MMP-13) with peptides in the active site defines exosites in the catalytic domain. FASEB J. 2013;27(11):4395–4405. doi:10.1096/fj.13-233601

50. Gomis-Rüth FX, Gohlke U, Betz M, et al. The Helping Hand of Collagenase-3 (MMP-13): 2.7 Å Crystal Structure of its C-terminal Haemopexin-like Domain. J Mol Biol. 1996;264(3):556–566. doi:10.1006/jmbi.1996.0661

51. Chung L, Dinakarpandian D, Yoshida N, et al. Collagenase unwinds triple-helical collagen prior to peptide bond hydrolysis. EMBO J. 2004;23(15):3020–3030. doi:10.1038/sj.emboj.7600318

52. Lauer-Fields JL, Tuzinski KA, Shimokawa K ichi, Nagase H, Fields GB. Hydrolysis of Triple-helical Collagen Peptide Models by Matrix Metalloproteinases. J Biol Chem. 2000;275(18):13282–13290. doi:10.1074/jbc.275.18.13282

53. Serwanja J, Wieland AC, Haubenhofer A, Brandstetter H, Schönauer E. A conserved strategy to attack collagen: The activator domain in bacterial collagenases unwinds triple-helical collagen. Proc Natl Acad Sci. 2024;121(16):e2321002121. doi:10.1073/pnas.2321002121

54. Manka SW, Carafoli F, Visse R, et al. Structural insights into triple-helical collagen cleavage by matrix metalloproteinase 1. Proc Natl Acad Sci. 2012;109(31):12461–12466. doi:10.1073/pnas.1204991109

55. Arnold LH, Butt LE, Prior SH, Read CM, Fields GB, Pickford AR. The Interface between Catalytic and Hemopexin Domains in Matrix Metalloproteinase-1 Conceals a Collagen Binding Exosite. J Biol Chem. 2011;286(52):45073–45082. doi:10.1074/jbc.M111.285213

56. Little CB, Barai A, Burkhardt D, et al. Matrix metalloproteinase 13–deficient mice are resistant to osteoarthritic cartilage erosion but not chondrocyte hypertrophy or osteophyte development. Arthritis Rheum. 2009;60(12):3723–3733. doi:10.1002/art.25002

57. Neuhold LA, Killar L, Zhao W, et al. Postnatal expression in hyaline cartilage of constitutively active human collagenase-3 (MMP-13) induces osteoarthritis in mice. J Clin Invest. 2001;107(1):35–44. doi:10.1172/JCI10564

58. Mehana ESE, Khafaga AF, El-Blehi SS. The role of matrix metalloproteinases in osteoarthritis pathogenesis: An updated review. Life Sci. 2019;234:116786. doi:10.1016/j.lfs.2019.116786

59. Hu Q, Ecker M. Overview of MMP-13 as a Promising Target for the Treatment of Osteoarthritis. Int J Mol Sci. 2021;22(4):1742. doi:10.3390/ijms22041742

60. Fingleton B. Matrix Metalloproteinases as Valid Clinical Target. Curr Pharm Des. 2007;13(3):333–346. doi:10.2174/138161207779313551

61. Nara H, Sato K, Kaieda A, et al. Design, synthesis, and biological activity of novel, potent, and highly selective fused pyrimidine-2-carboxamide-4-one-based matrix metalloproteinase (MMP)-13 zinc-binding inhibitors. Bioorg Med Chem. 2016;24(23):6149–6165. doi:10.1016/j.bmc.2016.09.009

62. Engel CK, Pirard B, Schimanski S, et al. Structural Basis for the Highly Selective Inhibition of MMP-13.

63. De Savi C, Waterson D, Pape A, et al. Hydantoin based inhibitors of MMP13—Discovery of AZD6605. Bioorg Med Chem Lett. 2013;23(16):4705–4712. doi:10.1016/j.bmcl.2013.05.089

64. Zhang F, Lee K, Linhardt R. SPR Biosensor Probing the Interactions between TIMP-3 and Heparin/GAGs. Biosensors. 2015;5(3):500–512. doi:10.3390/bios5030500

