## Supplementary figures and images for "Heparan sulfate selectively inhibits the collagenase activity of matrix metalloproteinase 13"

### Supplemental Figure 1

#### Supplemental Figure 1

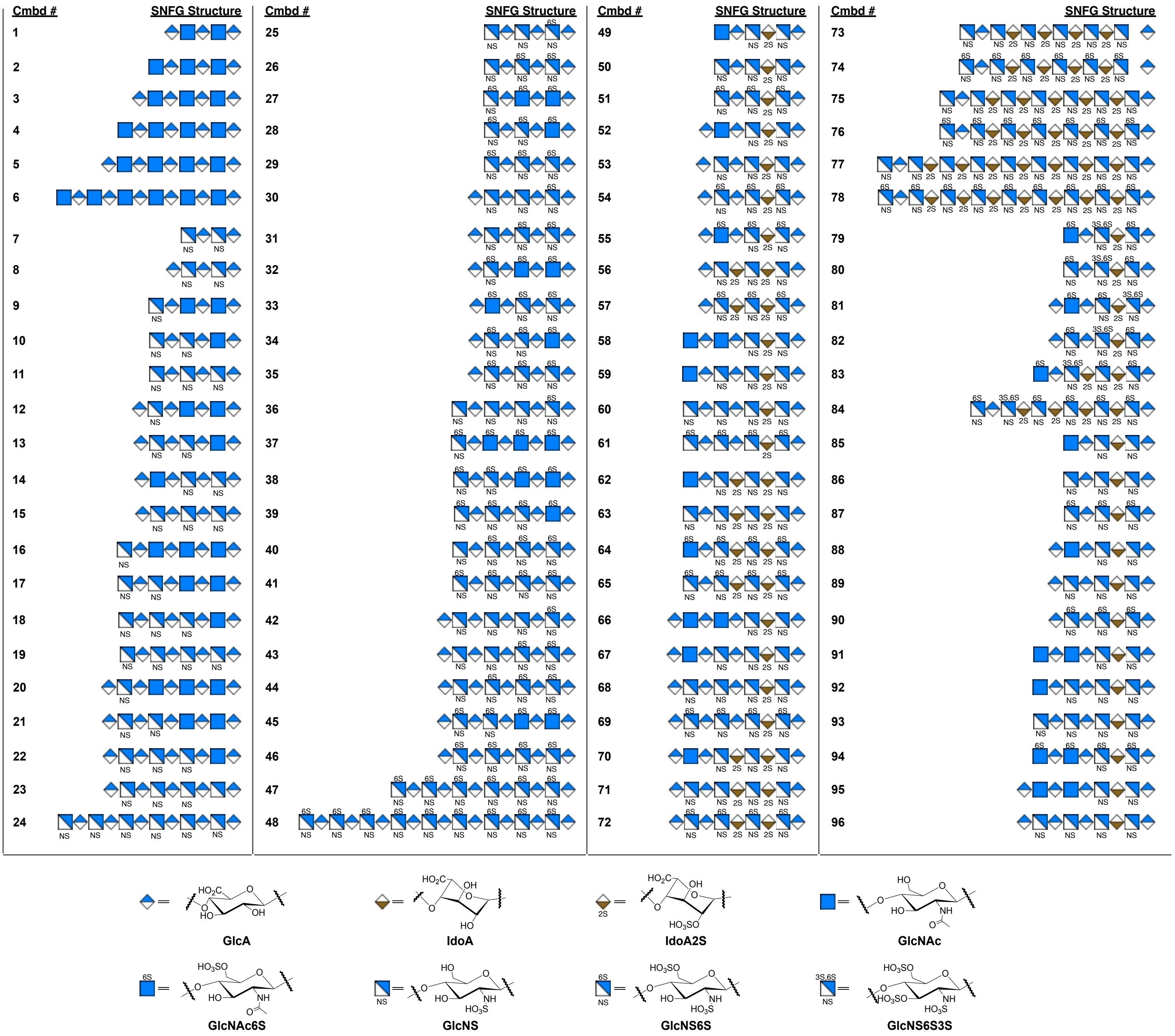
